# Chemical validation of *Mycobacterium tuberculosis* phosphopantetheine adenylyltransferase using fragment linking and CRISPR interference

**DOI:** 10.1101/2020.09.04.280388

**Authors:** Jamal El Bakali, Michal Blaszczyk, Joanna C. Evans, Jennifer A. Boland, William J. McCarthy, Marcio V. B. Dias, Anthony G. Coyne, Valerie Mizrahi, Tom L. Blundell, Chris Abell, Christina Spry

**Affiliations:** Department of Chemistry, University of Cambridge, Lensfield Road, Cambridge CB2 1EW (UK); Department of Biochemistry, University of Cambridge, 80 Tennis Court Road, Cambridge CB2 1GA (UK); MRC/NHLS/UCT Molecular Mycobacteriology Research Unit, DST/NRF Centre of Excellence for Biomedical TB Research & Wellcome Centre for Infectious Diseases Research in Africa, Institute of Infectious Disease and Molecular Medicine and Department of Pathology, Faculty of Health Sciences, University of Cape Town, Anzio Road, Observatory 7925 (South Africa)

**Keywords:** Coenzyme A, Drug discovery, Enzymes, Fragment-based, Tuberculosis

## Abstract

The coenzyme A (CoA) biosynthesis pathway has attracted attention as a potential target for much-needed novel antimicrobial drugs, including for the treatment of tuberculosis (TB), the lethal disease caused by *Mycobacterium tuberculosis (Mtb*). Seeking to identify the first inhibitors of *Mtb* phosphopantetheine adenylyltransferase (*Mtb*PPAT), the enzyme that catalyses the penultimate step in CoA biosynthesis, we performed a fragment screen. In doing so, we discovered three series of fragments that occupy distinct regions of the *Mtb*PPAT active site, presenting a unique opportunity for fragment linking. Here we show how, guided by X-ray crystal structures, we could link weakly-binding fragments to produce an active site binder with a K_D_ < 20 μM and on-target anti-*Mtb* activity, as demonstrated using CRISPR interference. This study represents a big step toward validating *Mtb*PPAT as a potential drug target and designing a *Mtb*PPAT-targeting anti-TB drug.

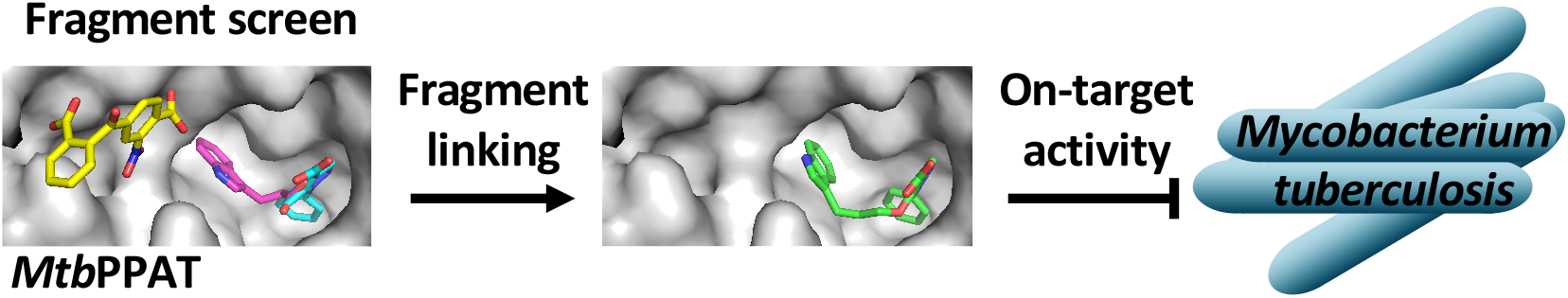

---

*Mycobacterium tuberculosis* (*Mtb*) is the causative agent of tuberculosis (TB), an infectious disease that predominantly affects the lungs and which was responsible for 1.5 million deaths in 2018 (including 0.3 million among people with HIV).^[1]^ The recommended treatment for drug-susceptible TB is a 6-month regimen of a combination of four first-line drugs, namely isoniazid, rifampicin, pyrazinamide and ethambutol.^[1-2]^ Poor adherence to TB therapies has contributed to the emergence of multidrug-resistant TB (MDR-TB), which poses a major public health threat.^[1]^ MDR-TB requires long-term treatment with second-line drugs that have limited efficacy and are often associated with considerable side effects.^[1]^ Therefore, there is an urgent need for new treatments with novel modes of action that will evade pre-existing resistance mechanisms.

Sequencing of the complete *Mtb* genome facilitated the identification of a number of potential new targets.^[3]^ On the basis of gene essentiality studies, the coenzyme A (CoA) biosynthetic pathway has been identified as a prospective target^[4]^. The synthesis of CoA has also very recently been implicated as a target of the anti-TB drug pyrazinamide.^[5]^ CoA is a vital cofactor that functions as an acyl group carrier or carbonyl-activating group in several essential biochemical transformations. Generally, CoA is synthesised from pantothenate (vitamin B_5_) in five steps.^[6]^ Phosphopantetheine adenylyltransferase (PPAT), a hexameric protein encoded by the *coaD* gene, catalyses the penultimate step – the transfer of an adenylyl group from ATP to 4’-phosphopantetheine, to form dephospho-CoA and pyrophosphate. Despite its previous classification as a non-essential gene based on transposon mutagenesis studies^[4b, 4d]^, a targeted gene disruption study provided evidence that the *coaD* gene is, in fact, essential for growth of *Mtb in vitro*.^[7]^ Moreover, little sequence identity^[8]^ and structural similarity is shared between *Mtb*PPAT and the human counterpart and as such the enzyme is considered an attractive target for anti-TB drug discovery.

Although no inhibitors of *Mtb*PPAT have been reported so far, high-throughput screens have led to the identification of two series of bacterial PPAT inhibitors.^[9]^ Furthermore, compounds from one of the series, have been shown to inhibit growth of *Staphylococcus aureus* and *Streptococcus pneumoniae in vitro* through inhibition of PPAT, and display some effect in two mouse efficacy models, thereby validating PPAT as a novel antimicrobial target.^[9a]^ Additionally, recent publications from Novartis describe the discovery of a series of inhibitors of *Escherichia coli* and *Pseudomonas aeruginosa* PPAT that were selective for these bacterial PPATs over their human orthologue.^[10]^ Herein, we describe a fragment-based approach^[11]^ leading to the discovery of the first non-natural *Mtb*PPAT ligands. Furthermore, we show, using a PPAT conditional knockdown strain, that the lead molecule possesses on-target anti-proliferative activity against *Mtb in vitro*, thereby chemically validating *Mtb*PPAT as a potential anti-TB drug target.

To identify starting points for inhibitor design, we screened a library of 1265 rule-of-three compliant^[12]^ fragments using a screening cascade combining orthogonal biophysical techniques.^[13]^ Fragments were initially screened using a fluorescence-based thermal shift assay (hits presented in Figure S1). Subsequently, saturation transfer difference (STD) and water-ligand observed via gradient spectroscopy (WaterLOGSY) NMR were used for hit confirmation and to identify active site binders (i.e. fragments binding competitively with active site binders ATP and CoA). The fragment screen (summarised in Figure 1) yielded three chemically-distinct fragment hits: benzophenone **1**, indole **2** and pyrazole **3**. Additional *Mtb*PPAT active site binders (e.g. **4-18**, Figure 1) were identified by testing analogues of each of the fragment hits using the thermal shift assay and/or STD and WaterLOGSY NMR (Figure S2-S4). Quantitative measurement of the fragments’ affinity for *Mtb*PPAT was subsequently attempted using isothermal titration calorimetry (ITC). Although heats of binding were detected upon titration of *Mtb*PPAT (24 μM, hexamer concentration) with some benzophenone and pyrazole fragments (at 10 mM), the binding isotherms showed poor signal-to-noise ratios, making it difficult to fit the data to a binding model reliably. This is likely a consequence of the invariably low affinity of fragments for their target^[11a]^ and/or a low enthalpic contribution to binding as has been observed for ATP binding.^[14]^

**Figure 1.**
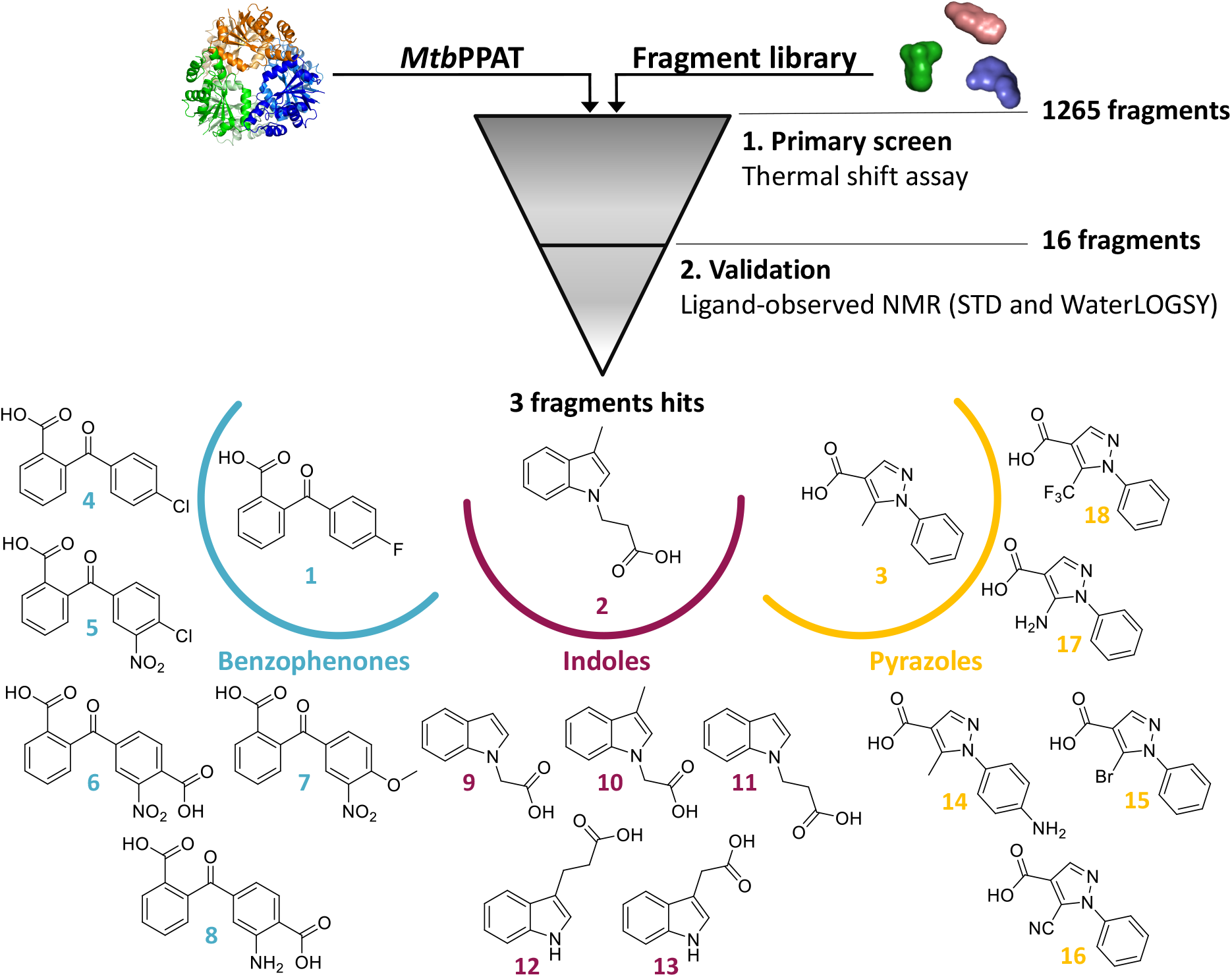
Screening cascade leading to the identification of 3 different fragment hits against *Mtb*PPAT. Analogues of each fragment hit subsequently found to bind *Mtb*PPAT are also shown.

By contrast with the fragment hits, a well-defined binding isotherm was observed upon titration of *Mtb*PPAT with CoA (Figure S5A), consistent with previous findings.^[14]^ Under near-identical conditions to those used by others to study ligand binding to *Mtb*PPAT by ITC,^[14]^ a slight deviation from a sigmoidal binding isotherm was observed. This is consistent with the previously reported asymmetry in the *Mtb*PPAT quaternary structure.^[14]^ Nonetheless, the data were a reasonable fit for a “one set of sites” independent binding model, from which a dissociation constant (K_D_) of 2.9 μM was estimated for CoA, a value only slightly lower than previous estimates (≥ 13 μM^[14]^). In turn, using CoA as a competitive binder, it was possible to estimate the affinity of indole series fragments in competition ITC experiments (Figure S5B), with a K_D_ of 0.9 mM determined for indole fragment **2** (ligand efficiency,^[15]^ LE = – ΔG (kcal.mol^−1^) per non-hydrogen atom (NHA) = 0.28). However, analogous experiments performed with benzophenone and pyrazole fragments produced clearly non-symmetrical binding isotherms (e.g. Figure S5C), indicative of the active sites of hexameric *Mtb*PPAT not being independent under these conditions. This complicated the assessment of affinity of the pyrazole and benzophenone fragments by competition ITC. Pyrazole series fragments were instead further evaluated by competition-based STD-NMR using the original pyrazole fragment **3** as a reporter molecule.^[16]^ This was possible due to the distinctive ^1^H signal of the C5-methyl group unique to this fragment (and fragment **14**, which was therefore excluded from the analysis). The analogues of pyrazole **3** all showed competition for the fragment’s binding site, as evidenced by the decreased STD signal of **3**’s C5-methyl group, with the following affinity for the site inferred from the signals observed: **15** > **16** > **18** > **17** (Figure S6).

To elucidate the binding modes of the fragment hits, crystals of *apo MtbPPAT* (protomer structure shown in Figure 2A) were individually soaked with fragments from each of the three series. Cocrystal structures (Table S1) demonstrated that the benzophenone, indole and pyrazole fragments bind to distinct positions within the *Mtb*PPAT active site, and, by contrast with a recently reported fragment screen targeting *E. coli* PPAT,^[10a]^ in both the phosphopantetheine and ATP binding sites. Benzophenone fragment **6** was observed to bind in the phosphopantetheine binding site while the indole and pyrazole fragments occupy the ATP binding site (Figure 2B). Benzophenone **6** was observed to bind *Mtb*PPAT with one carboxylic acid moiety oriented towards the back of the phosphopantetheine binding pocket, away from the solvent channel, forming a hydrogen bond with Val73, and interacting via water molecules (not shown) with Gly71, Val74 and Asn105 (Figure 2C). With the exception of the water-bridged hydrogen bond to Gly71, these interactions are fulfilled by the distal amide carbonyl of the enzyme’s natural substrate phosphopantetheine.^[14, 17]^ The phenyl ring is also positioned to form hydrophobic interactions with the sulfur of Met101. The 3-nitro substituent on the second phenyl ring forms an electrostatic interaction with the side chain of Lys87 (a residue that forms a water-mediated hydrogen bond with phosphopantetheine), and the carboxylic acid substituent forms a hydrogen bond with Gly8. Surprisingly, despite occupying a site distal to the ATP binding pocket, benzophenone **6** induces a similar conformational change to that observed upon ATP binding.^[14]^ This change predominantly affects Leu89-Arg90-Thr91-Gly92-Thr93-Asp94 (Figure 2D) and extends α-helix α4 by four residues.

**Figure 2.**
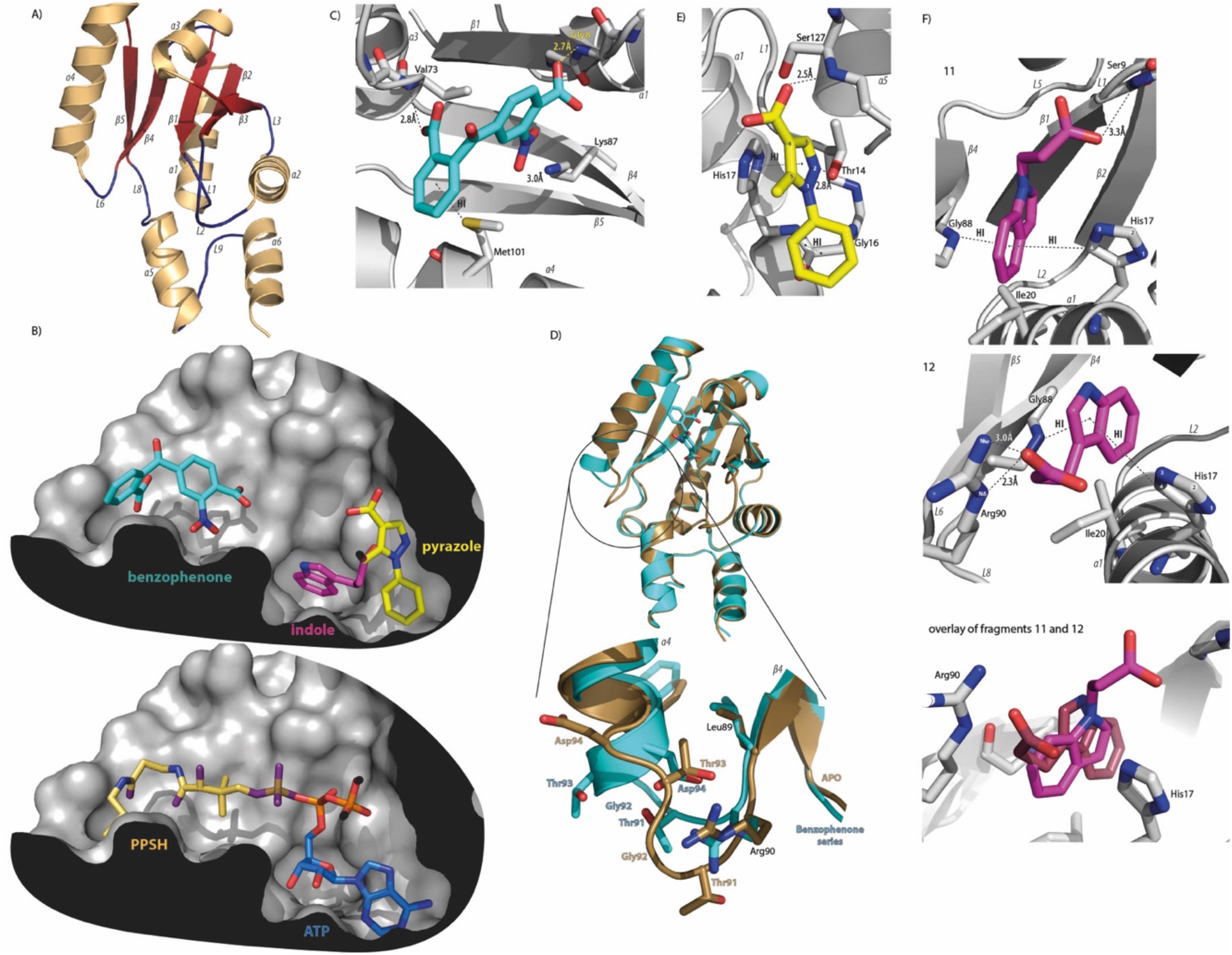
X-ray crystal structures of *Mtb*PPAT with and without fragments bound. A) *apo Mtb*PPAT protomer shaded according to secondary structure: gold, α-helix; red, β-strand; blue, loop). B) Binding positions of benzophenone, pyrazole and indole fragment hits **6**, **3** and **12**, respectively, in the active site. Binding poses of the natural substrates phosphopantetheine (PPSH) and ATP are shown for comparison. C) Hydrogen-bonding interactions and hydrophobic interactions (HI) (both represented by dashed lines) between benzophenone **6** and *Mtb*PPAT. D) Conformational changes observed upon binding of benzophenone **6** to *Mtb*PPAT. The *apo Mtb*PPAT structure is shown in gold and the fragment-bound structure in cyan. The zoomed-in image displays the region of *Mtb*PPAT that is most affected, with the side chains of Leu89, Arg90, Thr91, Gly92, Thr93 and Asp94 shown. E) Hydrogen-bonding and hydrophobic interactions between pyrazole **3** and *Mtb*PPAT. F) Hydrogen-bonding and hydrophobic interactions between indoles **11** and **12** and *Mtb*PPAT. An overlay of fragments **11** and **12** in the pocket is also shown for reference. In all structures, oxygen atoms are shown in red or, in the case of PPSH, purple, and nitrogen atoms are shown in blue. Protein Data Bank entries: 6QMI (**3**), 6G6V (**6**), 6QMF (**11**), 6QMG (**12**). In C, E and F, only direct (not water-mediated) hydrogen-bonding interactions are shown.

Binding of pyrazole **3** to *Mtb*PPAT induces a conformational change akin to that observed upon benzophenone **6** and ATP binding. As shown in Figure 2B, the phenyl substituent of pyrazole **3** binds in the position occupied by the adenine ring of ATP and the carboxylic acid binds in the position of the β-and γ-phosphates. Thr14 and Ser127, two residues that interact directly, or via a water molecule, with ATP, form hydrogen bonds with the pyrazole N2 nitrogen and the carboxylic acid, respectively (Figure 2E). Two conserved water molecules in the active site (held in a network by Phe10 and His17) are seen to form an additional interaction with the carboxylic acid group of this fragment (not shown); this same interaction is fulfilled by the α-phosphate of ATP. Additionally, the phenyl group of the bound fragment is involved in a π-π interaction with the Gly16-His17 peptide bond, and the pyrazole ring is involved in a π-π stacking interaction with His17.

Indole fragments **11** and **12** were observed to bind *Mtb*PPAT in overlapping positions, but to differ in their orientation (Figure 2F). The indole moieties of the fragments bind near to the site that the ribose of ATP occupies, and in the case of **12**, extends into a small pocket behind the ATP binding site. The carboxylic acid is positioned either near to where the α-phosphate of ATP binds (**11**), forming hydrogen bonds with His17 and Ser9 (either directly or via water molecules), or between His17 and Arg90 (**12**), forming a hydrogen bond with the latter residue. His17, Ser9 and Arg90 are all residues involved in ATP binding.

Pyrazole fragment **3** and indole fragment **12** have subsequently been co-crystallised with the closely-related PPAT from *Mycobacterium abcessus* (79% identical, 87% similar).^[18]^ Interestingly, the observed binding mode of the pyrazole fragment is identical. However, by contrast, the indole is found in the phosphopantetheine binding site, even though another indole (fragment **2**) co-crystallises with *Mab*PPAT in a site corresponding to the fragment **12** binding site in *Mtb*PPAT.

With fragments binding to distinct regions of the *Mt*bPPAT active site, there were multiple opportunities for fragment elaboration, including fragment linking. The present work focuses on linking of the indole and pyrazole fragments that bind in close proximity. Of the indole fragments observed to bind *Mtb*PPAT, indole **12** (K_D_ = 1.0 mM, LE = 0.29; Figure S7) was selected, in part, because the alkyl carboxylic acid substituent at the C-5 position points towards the C-3 position of pyrazole **3** (Figure 2B). Elaborated compounds with flexible linkers predicted to allow the two parent fragments to bind in their preferred position were designed, and because of chemical tractability, synthesis of ether-based linkers was favoured. Three compounds (**19** – **21**) with ether linkers of varying lengths (Figure 3A) were synthesised as described in Scheme S1 and tested against *Mtb*PPAT. All three compounds were observed to bind *Mtb*PPAT by NMR and to do so competitively with CoA (e.g. Figure S8). Furthermore, binding to *Mtb*PPAT was detectable, directly, by ITC. A K_D_ of ~ 40 μM (~ 25-fold lower than the K_D_ determined for the parent indole fragment **12**) was determined for compound **19** (with a three-unit linker; Figure S9) and a further increase in affinity was observed upon extending the linker by one or two carbons. K_D_ values of 15 ± 4 and 27 ± 4 μM were determined for compound **20** (with a four unit linker) and **21** (with a five unit linker) under the same conditions, corresponding to LE values of 0.24 and 0.22 kcal.mol^−1^.NHA^−1^, respectively; Figures 3A and S10-11). In the case of **20**, this corresponds to an affinity increase of ~65-fold relative to the parent indole fragment. An X-ray co-crystal structure of compound **20** bound to *Mtb*PPAT was obtained by soaking *Mtb*PPAT crystals with the compound. The structure revealed that as intended, the 4-unit ether linker allowed the indole and pyrazole moieties to bind in the same sites as the parent fragments. However, although the pose of the pyrazole moiety is almost identical to that of the parent pyrazole, the indole moiety is slightly tilted compared to the indole moiety of the parent fragment (Figure 3B). Compound **20** is involved in multiple polar interactions (Figure 3C). The –NH-of the indole ring makes a hydrogen bond with the carbonyl of Pro7, an interaction that was not seen with the parent fragment, while the ether linker interacts with Arg90. The pyrazole ring of **20** is involved in an aromatic interaction with His17 and a hydrogen bond with Thr14 via Nitrogen N2. The carboxylic acid forms hydrogen bonds with Ser127 and a water-mediated hydrogen bond with Ser128.

**Figure 3.**
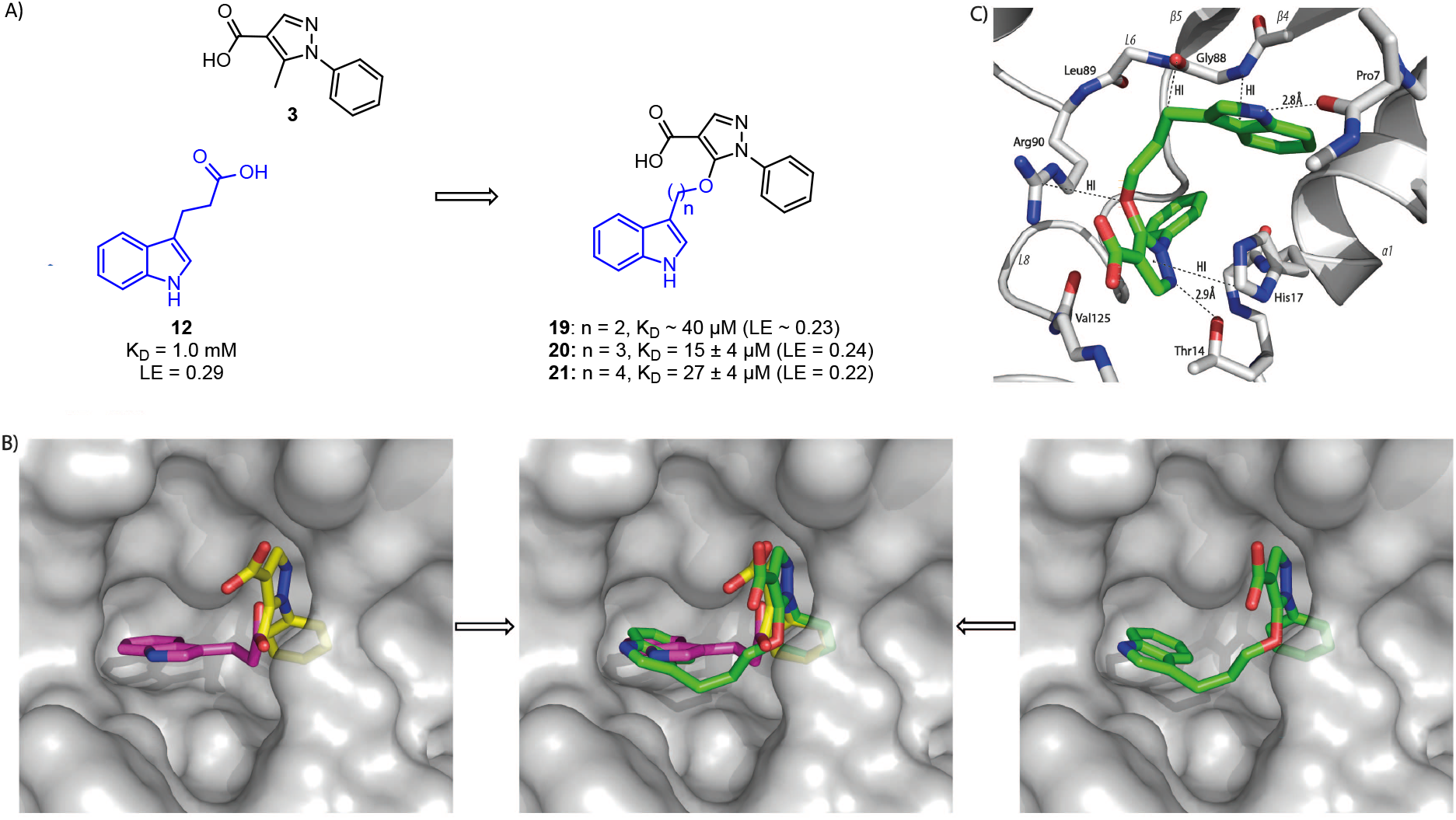
Fragment elaboration leading to compounds **19 – 21**. A) Chemical structures of the parent fragments and linked compounds synthesized, with K_D_ values and LEs (in kcal.mol^−1^.NHA^−1^) determined shown. B) Overlay of fragment **3** (carbons in yellow) and fragment **12** (carbons in magenta) in the active site of *MtbPPAT* (LHS), the co-crystal structures of compound **20** (carbons in green) bound to *Mtb*PPAT (RHS), and an overlay of all three molecules in the active site (centre). C) Hydrogen-bonding interactions and hydrophobic interactions (HI) (both represented by dashed lines) between compound **20** and *Mtb*PPAT. For clarity, only direct hydrogen-bonding interactions are shown and the orientation of compound **20** differs to that shown in B. The hydrogen-bond between the carboxylic acid of **20** and Ser127 is not shown. Protein Data Bank entries: 6QMI (**3**), 6G6V (**6**), 6QMH (**20**).

Despite keeping a binding mode very close to the parent fragments, the linking strategy did not lead to additivity or super-additivity as could be expected in such a case^[19]^. This is likely due to the high flexibility of the 4-unit ether linker. Moving forward, it may be possible to improve affinity by rigidifying this linker in order to reduce the entropic cost of binding.

*Mtb* hypomorphs that conditionally underexpress genes of interest have successfully been utilised to demonstrate on-target whole-cell activity.^[20]^ In the case of *coaD*, however, it has been difficult to achieve a reduction in transcript using the traditional approach of promoter-replacement, due to the low basal level of expression of this gene in wild-type *Mtb*.^[4a]^ We therefore employed CRISPR interference (CRISPRi)-mediated transcriptional silencing^[21]^ to investigate whether **20** inhibits proliferation of *Mtb* H37RvMA^[22]^ via inhibition of PPAT activity. The utility of CRISPRi for validation of compound mode of action against *Mtb in vitro* was recently demonstrated.^[23]^ Using an analogous approach, a *coaD*-targeting single guide RNA (sgRNA) was identified that, when coexpressed in *Mtb* H37RvMA with a “nuclease-dead” Cas9 (dCas9)^[21]^ (both from anhydrotetracycline (ATc)-inducible promoters), mediated inhibition of *Mtb* H37RvMA proliferation in the presence of ATc (Figure S12, SI Experimental Methods). This provides independent confirmation of the previously reported essentiality of *coaD* for growth of *Mtb in vitro*^[7]^. This “*coaD* CRISPRi knockdown *Mtb* mutant” was used in checkerboard assays^[24]^ to assess the antibacterial activity of **20**. In the absence of ATc, compound **20** inhibited proliferation of the *coaD* CRISPRi *Mtb* mutant (minimum inhibitory concentration, MIC99 ~ 60 μM) and the effect was enhanced in an ATc concentration-dependent manner in the presence of 1.6 and 0.8 ng/mL ATc (the highest concentrations of ATc that permitted uninhibited proliferation in liquid culture in the absence of **20**) (Figure 4A). By contrast, ATc had no effect on the potency of **20** against wildtype *Mtb* H37RvMA (Figure 4B). Similarly, ATc had little-to-no effect on *Mtb* H37RvMA co-expressing dCas9 and a sgRNA targeting *coaBC* (the gene coding for the enzymes catalysing the previous two steps in the CoA biosynthesis pathway) (“*coaBC* CRISPRi knockdown *Mtb* mutant”; Figure S12) at the highest concentrations compatible with uninhibited proliferation in liquid culture in the absence of **20** (Figure 4C). Furthermore, ATc did not render the *coaD* CRISPRi *Mtb* mutant more susceptible to the standard TB drugs rifampicin and isoniazid (Figure 4D-E). These findings are consistent with *coaD* transcriptional silencing rendering *Mtb* H37RvMA selectively hypersusceptible to **20** and therefore with **20** inhibiting proliferation of *Mtb* by inhibiting the activity of PPAT.

**Figure 4.**
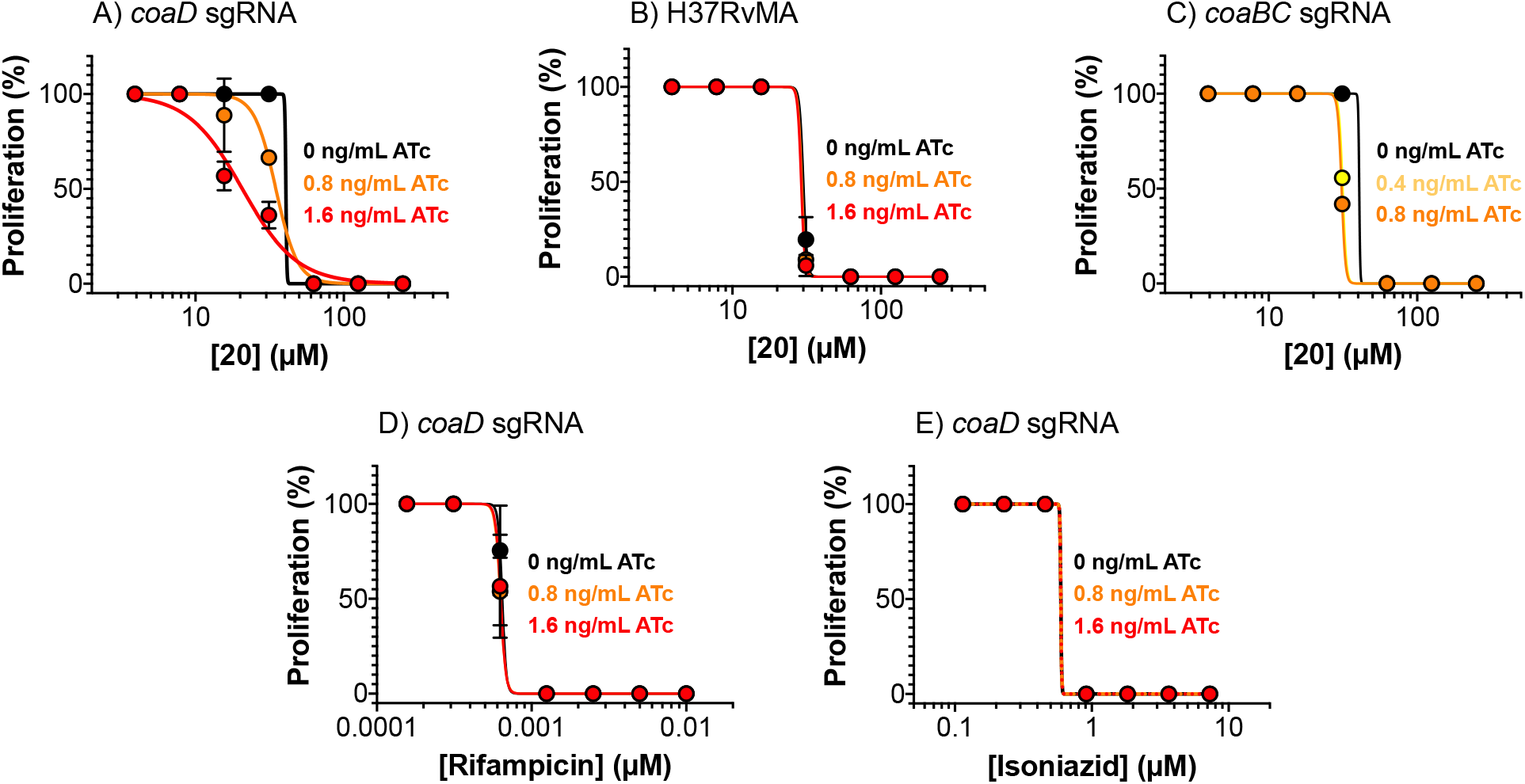
Transcriptional silencing of *coaD* renders *Mtb* hypersusceptible to **20.** Proliferation of *Mtb* H37RvMA co-expressing dCas9 and a *coaD*-targeting sgRNA (A, D-E), wild-type *Mtb* H37RvMA (B) or *Mtb* H37RvMA co-expressing dCas9 and a *coaBC*-targeting sgRNA (C) in the presence and absence of ATc and increasing concentrations of **20** (A-C), rifamipicin (D) or isoniazid (E) was monitored. The data are representative of two independent experiments. Averages of three replicates are shown and error bars represent standard deviation.

In conclusion, by using a fragment-based approach we have discovered the first low-micromolar non-natural *Mtb*PPAT active site binder that also displays on-target *in vitro* whole cell activity, as demonstrated using CRISPRi. The multiple fragment co-crystal structures obtained in this study were pivotal and facilitated fragment elaboration by linking, a fragment-elaboration strategy of which there are few successful examples in the literature.^[19]^ In addition to rigidifying the linker, future studies will focus on modifying both the phenyl and indole moieties to pick up additional interactions with surrounding amino acid residues and linking with the benzophenone fragment to ensure specificity of inhibition.

The combination of fragment-based and CRISPRi approaches used here has enabled chemical validation of *Mtb*PPAT. This not only provides justification for ongoing MtbPPAT-based drug discovery efforts but also provides additional evidence of the utility of target-based whole-cell screening (alone and in conjunction with fragment-based approaches) in TB drug discovery.

## Acknowledgements

We are grateful to DuPont for supplying several analogues of the initial fragment hits. We also thank Jeremy Rock and Sarah Fortune for providing the mycobacterial CRISPRi plasmids, and Peter Gimeson for advice on ITC analysis. The project was funded by grants from the Foundation for the National Institutes of Health with support from the Bill and Melinda Gates Foundation (grant #OPP1024021 to C.A. and T.L.B.), and the EU FP7 (project HEALTH-F3-2011-260872, More Medicines for Tuberculosis; to C.A. and T.L.B.). C.S. was funded, in part, by an NHMRC Overseas Biomedical Fellowship (1016357). J.C.E. was funded by a grant to V.M. from the Bill and Melinda Gates Foundation via a subaward from the Foundation for the National Institutes of Health (grant #OPP1158806), and with support from a Senior International Research Scholar Grant to V.M. from the Howard Hughes Medical Institute and grants from the South African Medical Research Council and National Research Foundation (to V.M).

## Notes

https://www.rcsb.org/structure/6QMI

https://www.rcsb.org/structure/6G6V

https://www.rcsb.org/structure/6QMF

https://www.rcsb.org/structure/6QMG

https://www.rcsb.org/structure/6QMH

### Competing Interest Statement

The authors have declared no competing interest.

