## Supplementary information for "Chemical validation of *Mycobacterium tuberculosis* phosphopantetheine adenylyltransferase using fragment linking and CRISPR interference"

Construction of CRISPRi conditional knockdown mutants…………………………..21

Effect of test compounds on *Mtb* growth……………………………………………..22

Synthetic organic chemistry…………………………………………………………..22

**References…………………………………………………………………………………31**

**NMR spectra………………………………………………………………………………32**

#### Supplementary Figures

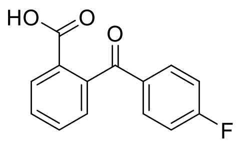

**1**

$\Delta T_m = +1.1 \pm 0.2$ °C

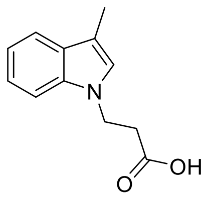

**2**

$\Delta T_m = +4.6 \pm 0.5$ °C

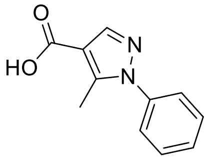

**3**

$\Delta T_m = +1.9 \pm 0.2$ °C

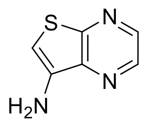

**S1**

$\Delta T_m = +0.9 \pm 0.1$ °C

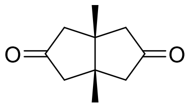

**S2**

$\Delta T_m = +2.2 \pm 0.5$ °C

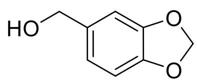

**S3**

$\Delta T_m = +1.0 \pm 0$ °C

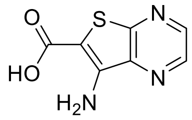

**S4**

$\Delta T_m = +0.8 \pm 0.2$ °C

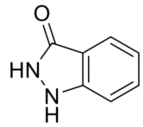

**S5**

$\Delta T_m = +1.1 \pm 0.1$ °C

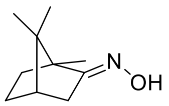

**S6**

$\Delta T_m = +1.9 \pm 0.2$ °C

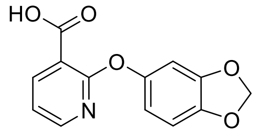

**S7**

$\Delta T_m = +1.0 \pm 0$ °C

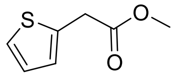

**S8**

$\Delta T_m = +0.8 \pm 0.2$ °C

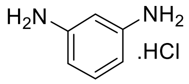

**S9**

$\Delta T_m = +1.0 \pm 0$ °C

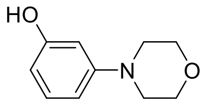

**S10**

$\Delta T_m = +1.0 \pm 0.1$ °C

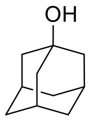

**S11**

$\Delta T_m = +2.4 \pm 0.1$ °C

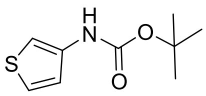

**S12**

$\Delta T_m = +1.0 \pm 0$ °C

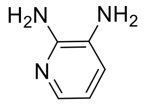

**S13**

$\Delta T_m = +0.8 \pm 0.1$ °C

**Figure S1:** Structures and thermal shift ($\Delta T_m$) values (in ˚C) of the fragment hits identified in a fluorescence-based thermal shift screen of 1265 fragments from a commercially available library (Maybridge). All fragments were tested at a concentration of 5 mM. Fragments shown, reproducibly (in two independent experiments) increased the melting temperature of *Mtb*PPAT by ≥ the 0.7 ˚C "hit" cut-off set. $\Delta T_m$ values shown are averages from two independent experiments and errors represent range/2.

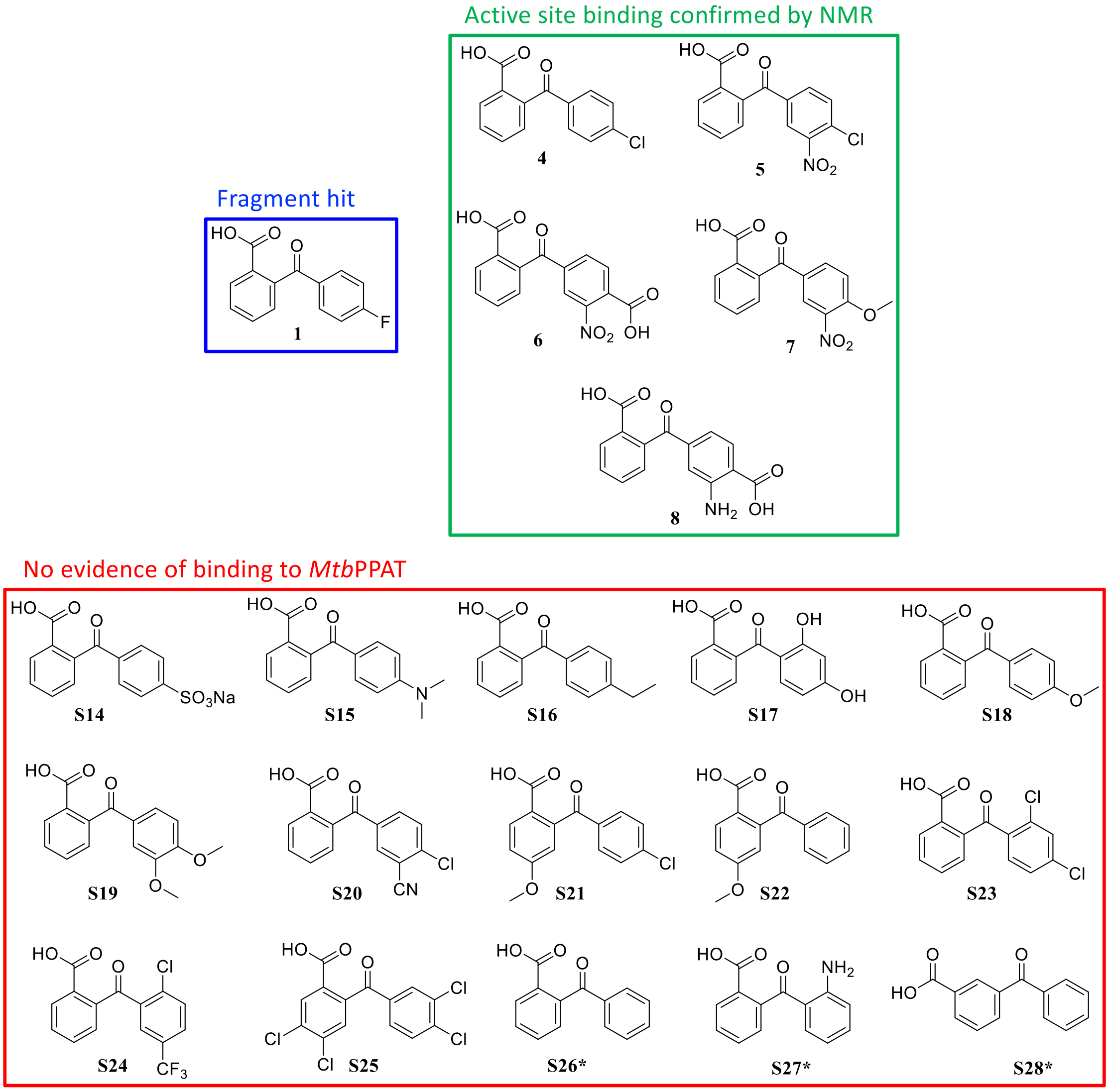

**Figure S2:** Analogues of fragment **1** tested for binding to *Mtb*PPAT. Analogues were obtained from DuPont (**6** – **8**, **S14** – **S25**) or purchased from commercial sources (**4**, **5** and **S26** – **S28**). Active site binding was confirmed using WaterLOGSY and STD NMR, with ligands present at 1 mM and *Mtb*PPAT at 20 µM; active site displacement was achieved using 1 mM ATP or CoA. *Fragments **S26** – **S28** were tested for active site binding by WaterLOGSY and STD NMR; other fragments without evidence of binding to *Mtb*PPAT were tested (at a concentration of 5 mM) by thermal shift assay only.

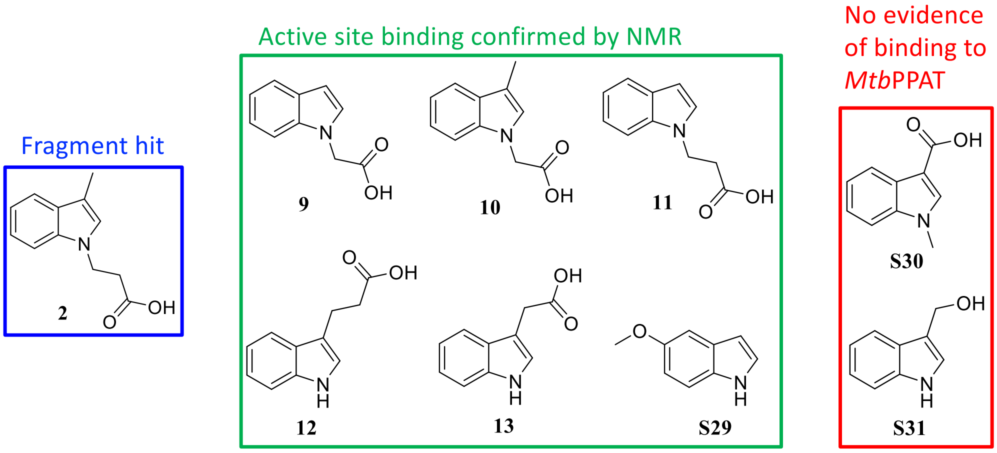

**Figure S3:** Analogues of **2** tested for binding to *Mtb*PPAT. Analogues **9-11** were synthesised according to Scheme S2, while analogues **12**, **13** and **S29 – 31** were purchased from commercial sources. Active site binding to *Mtb*PPAT was probed using ligand-based NMR experiments (WaterLOGSY and STD), with ligands present at 1 mM and *Mtb*PPAT at 20 µM; active site displacement was achieved using 1 mM ATP or CoA.

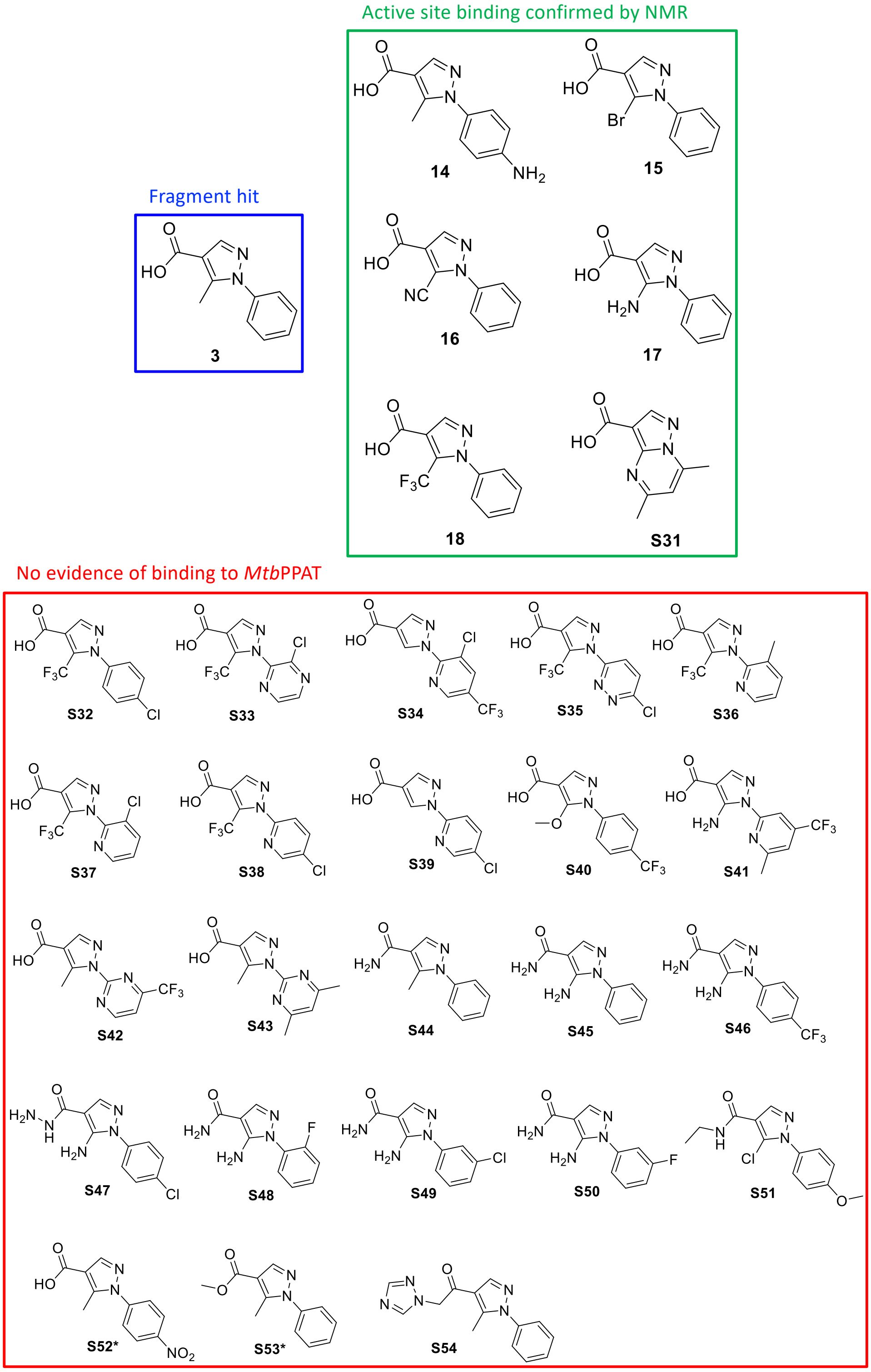

**Figure S4:** Analogues of fragment **3** tested for binding to *Mtb*PPAT. Analogues **14**, **S52** and **S53** were synthesised according to Scheme S3, while analogues **17** and **18** were purchased from commercial sources and analogues **15** – **16, S31** – **S51** and **S54** were obtained from DuPont. Active site binding was confirmed using ligand-based NMR experiments (WaterLOGSY and STD), with ligands present at 1 mM and *Mtb*PPAT at 20 µM; active site displacement was achieved using 1 mM ATP or CoA. *Fragments **S52** and **S53** were tested for active site binding by ligand-based NMR experiments (WaterLOGSY and STD); other fragments without evidence of binding to *Mtb*PPAT were tested (at a concentration of 5 mM) by thermal shift assay only.

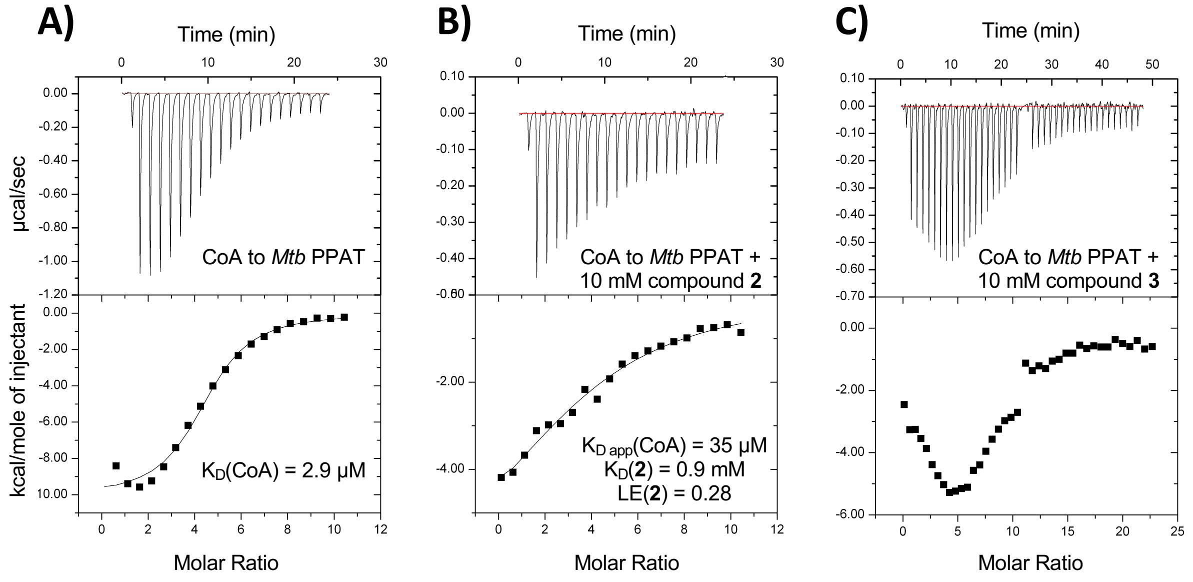

**Figure S5:** ITC analysis of *Mtb*PPAT binding to CoA, indole fragment **2** and pyrazole fragment **3**. Titration of *Mtb*PPAT with 600 µM CoA in the absence (**A**) or presence of 10 mM indole fragment **2** (**B**) or pyrazole fragment **3** (**C**). In **A** and **B**, the initial concentration of *Mtb*PPAT was 54 µM and in **C**, the initial concentration was 72 µM (protomer concentrations, equivalent to 9 and 12 µM hexamer, respectively). The upper panels show the change in energy required to maintain a constant temperature during the titration and the bottom panels show the integrated heats of binding. Titrations were performed in the presence of 30 mM HEPES, pH 8.0, 200 mM NaCl, 5 mM $MgCl_2$, 0.5 mM TCEP and 5% (v/v) DMSO. To the solution of *Mtb*PPAT, 19 × 2 µL test compound was injected. In **C**, where saturation of the protein was not achieved after 19 injections, a further 19 × 2 µL test compound was injected. Data in **A** and **B** are fitted to a "one set of sites" binding model (solid line). In **A** and **B**, binding stoichiometries (*n*) of 0.7 (relative to the protomer) were determined. The apparent $K_D$ ($K_{D\,app}$) determined for CoA in the presence of fragment **2** was used to calculate a $K_D$ for fragment **2** (as described in the experimental methods). The corresponding LE of indole fragment **2**, in $kcal.mol^{-1}.NHA^{-1}$, is shown.

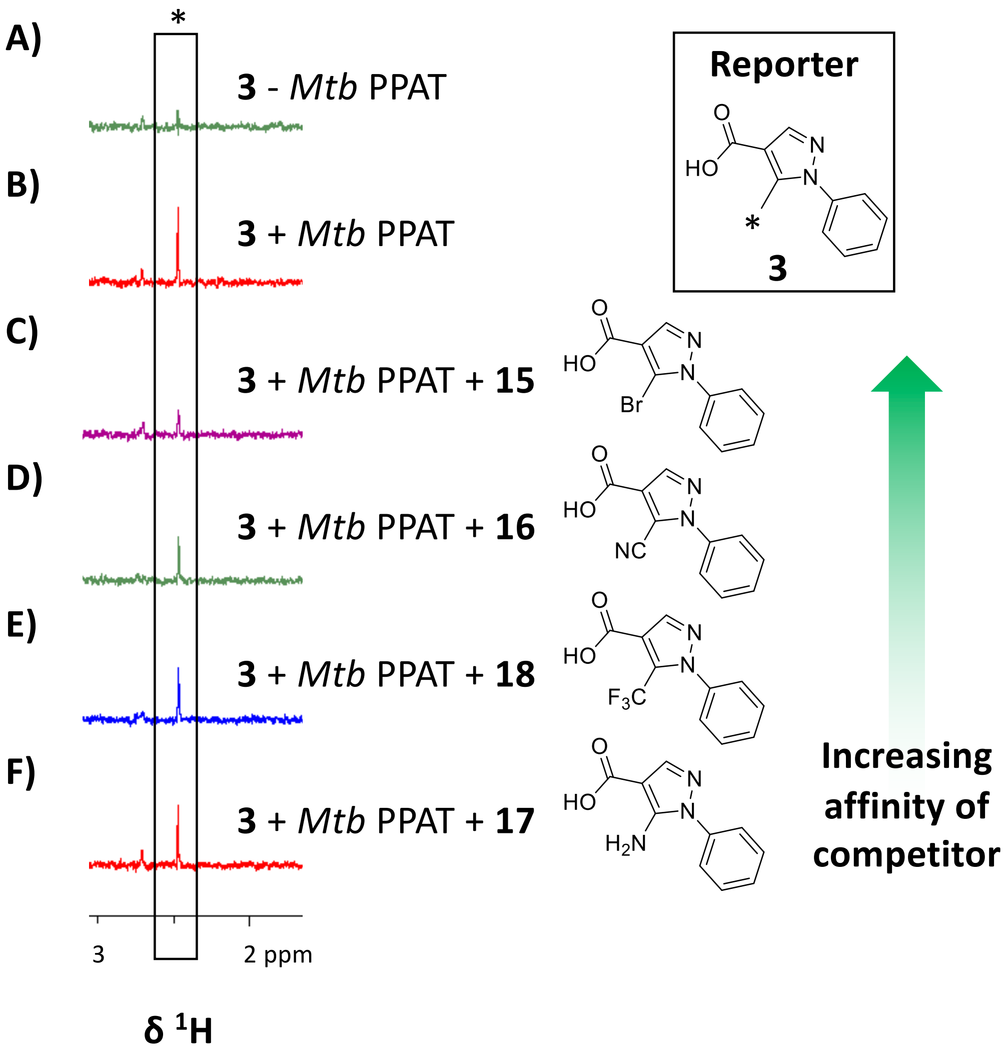

**Figure S6:** Ranking of pyrazole fragments based on affinity by competition-based STD-NMR. STD-NMR spectra showing the STD signal intensity of the C5-methyl protons (*) of pyrazole **3** (1 mM; the "reporter" molecule) in the absence (**A**) or presence (**B-F**) of 20 µM *Mtb*PPAT (protomer concentration, equivalent to 3.3 µM hexamer) and absence (**A-B**) or presence of 1 mM "competitor" fragments: pyrazole **15** (**C**), **16** (**D**), **18** (**E**) and **17** (**F**). The rank order of affinity inferred for the competitor molecules based on the STD signal of the reporter molecule, is shown.

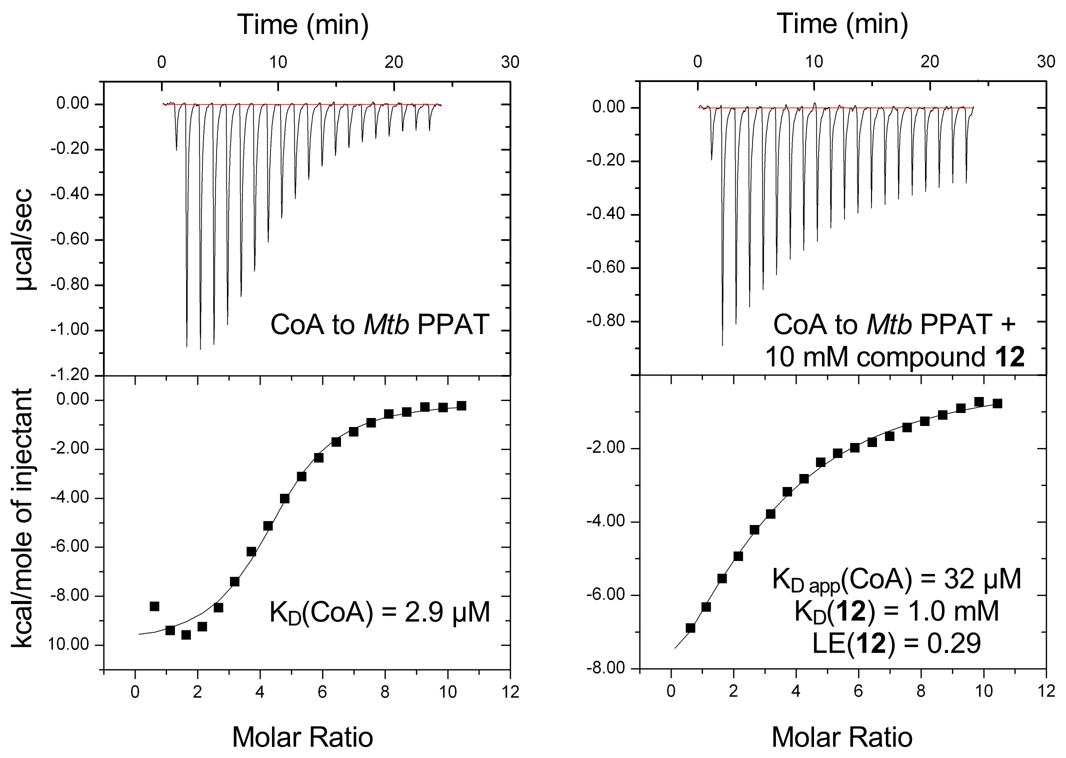

**Figure S7:** Competitive ITC analysis of *Mtb*PPAT binding to indole fragment **12**. Titration of *Mtb*PPAT with 600 µM CoA in the absence (LHS) or presence of 10 mM indole fragment **12** (RHS). The initial concentration of *Mtb*PPAT was 54 µM (protomer concentration, equivalent to 9 µM hexamer). The upper panels show the change in energy required to maintain a constant temperature during the titration and the bottom panels show the integrated heats of binding. Titrations were performed in the presence of 30 mM HEPES, pH 8.0, 200 mM NaCl, 5 mM $MgCl_2$, 0.5 mM TCEP and 5% (v/v) DMSO. To the solution of *Mtb*PPAT, 19 × 2 µL test compound was injected. Data are fitted to a "One Set of Sites" binding model (solid line). Binding stoichiometries (*n*) of 0.5 – 0.7 were determined. The LE of indole fragment **12**, in $kcal.mol^{-1}.NHA^{-1}$, is shown.

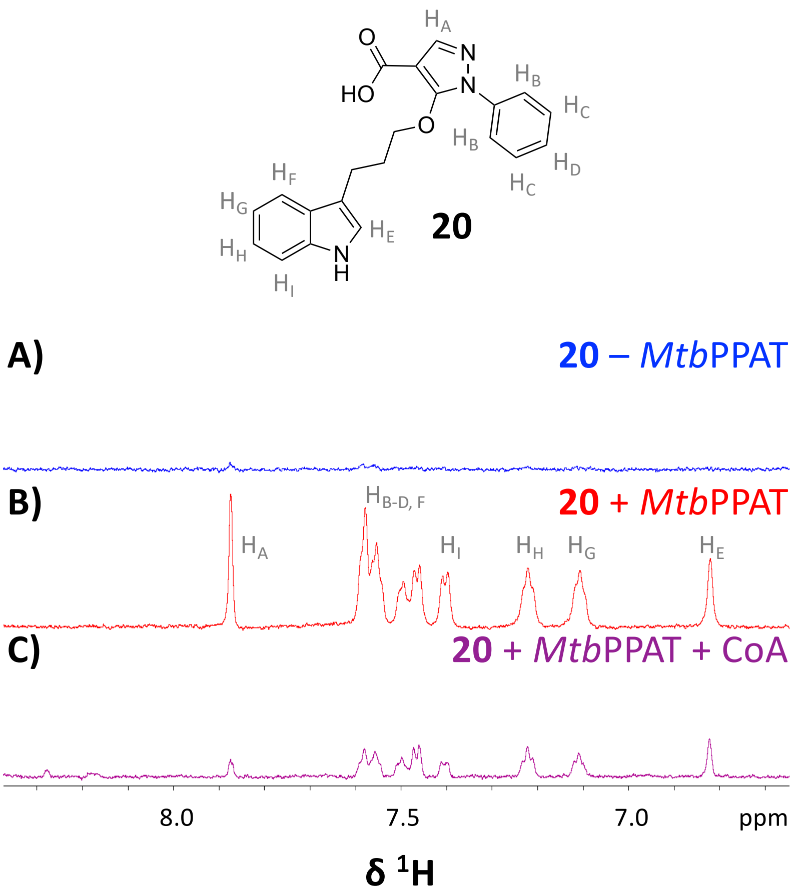

**Figure S8:** Detection of CoA-competitive binding of compound **20** to *Mtb*PPAT by STD-NMR. STD-NMR spectra showing the STD signals (or lack thereof) of the aromatic protons of compound **20** in the absence (**A**) or presence (**B-C**) of 20 µM *Mtb*PPAT (protomer concentration, equivalent to 3.3 µM hexamer) and absence (**A-B**) or presence of 1 mM CoA (**C**).

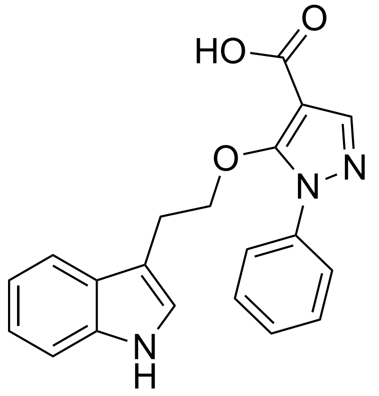

**19**

$K_D$ = 38 ± 2 μM
LE = 0.23

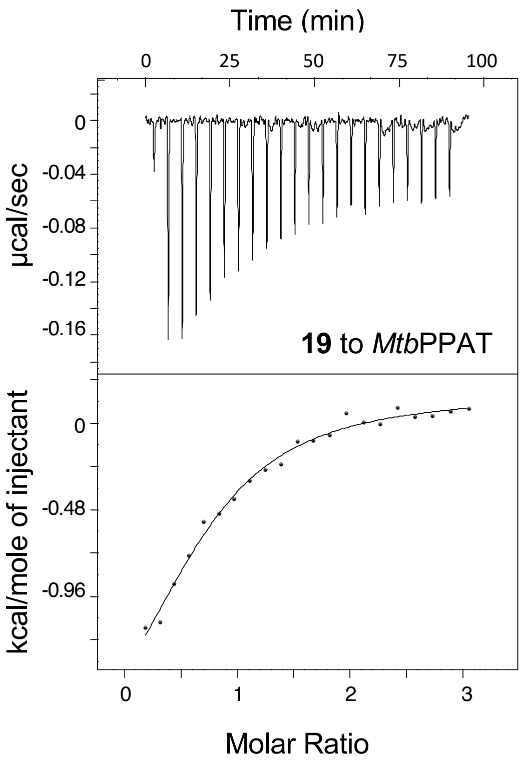

**Figure S9:** ITC analysis of *Mtb*PPAT binding to compound **19**. The upper panel shows the change in energy required to maintain a constant temperature during the titration of 100 μM *Mtb*PPAT (protomer concentration, equivalent to 17 μM hexamer) with compound **19** and the bottom panel shows the integrated heats of binding. Titrations were performed in the presence of 30 mM HEPES, pH 8.0, 20 mM NaCl, 0.5 mM TCEP and 5% (v/v) DMSO. To the solution of *Mtb*PPAT, 21 × 2 μL compound **19** (1 mM) was injected. Data are fitted to a "one set of sites" binding model (solid line). A binding stoichiometry (*n*) of 0.7 (relative to the protomer) was determined. The data are representative of two independent experiments and the average $K_D$ determined ± range/2 is shown. The corresponding LE, in kcal.mol$^{-1}$.NHA$^{-1}$, is also shown.

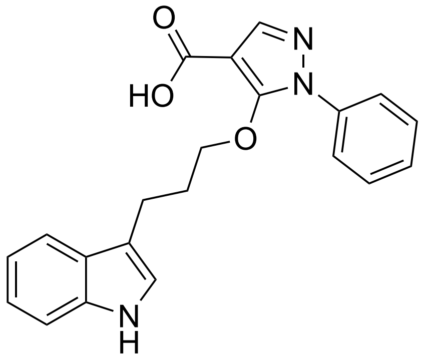

**20**

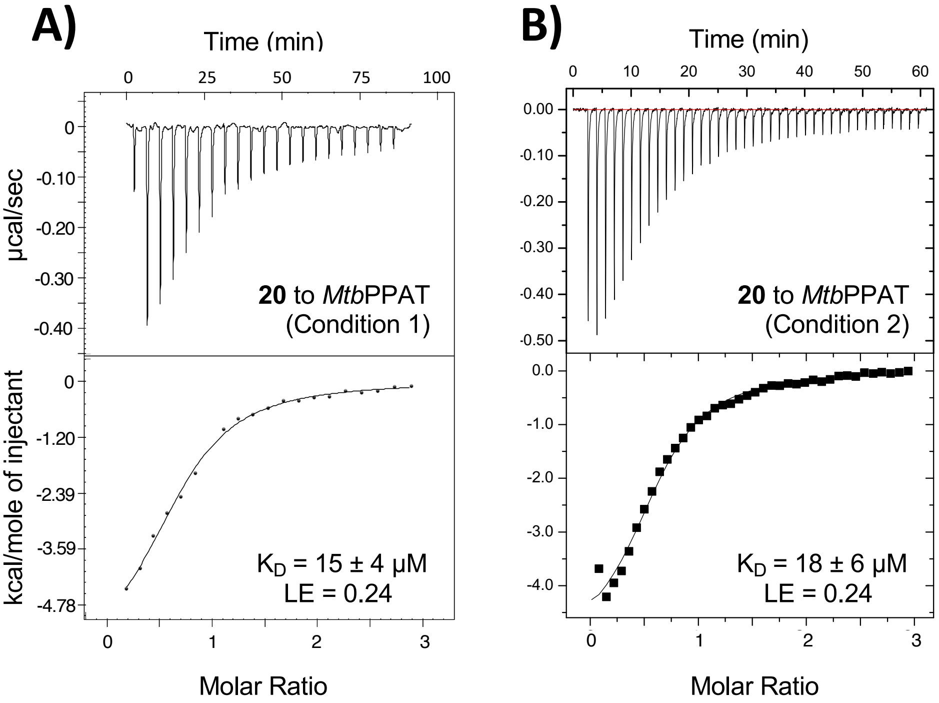

**Figure S10:** ITC analysis of *Mtb*PPAT binding to compound **20**. The upper panels show the change in energy required to maintain a constant temperature during the titration of *Mtb*PPAT with compound **20** and the bottom panels show the integrated heats of binding. Titrations were performed in the presence of 30 mM HEPES, pH 8.0, 20 mM NaCl, 0.5 mM TCEP and 5% (v/v) DMSO (**A**, "Condition 1", matching the conditions used to generate the data shown in Figures S9 and S11) or 30 mM HEPES, pH 8.0, 200 mM NaCl, 5 mM $MgCl_2$, 0.5 mM TCEP and 5% (v/v) DMSO (**B**, "Condition 2", matching the conditions used to generate the data shown in Figures S5 and S7). In **A**, 20 × 2 μL of 1 mM compound **20** was injected into a solution of 100 μM *Mtb*PPAT (protomer concentration, equivalent to 17 μM hexamer). In **B**, 39 × 1 μL of 1 mM compound **20** was injected into a solution of 72 μM *Mtb*PPAT (protomer concentration, equivalent to 12 μM hexamer). Data are fitted to a "one set of sites" binding model (solid line). Binding stoichiometries (*n*) of 0.6 (relative to the protomer) were determined. Each set of data shown is representative of two or three independent experiments. $K_D$ values shown are averaged from the two or three experiments and errors represent range/2 or standard error of the mean, respectively. The corresponding LEs, in $kcal.mol^{-1}.NHA^{-1}$, are also shown. Comparable $K_D$ (and LE) values were determined under the two sets of conditions.

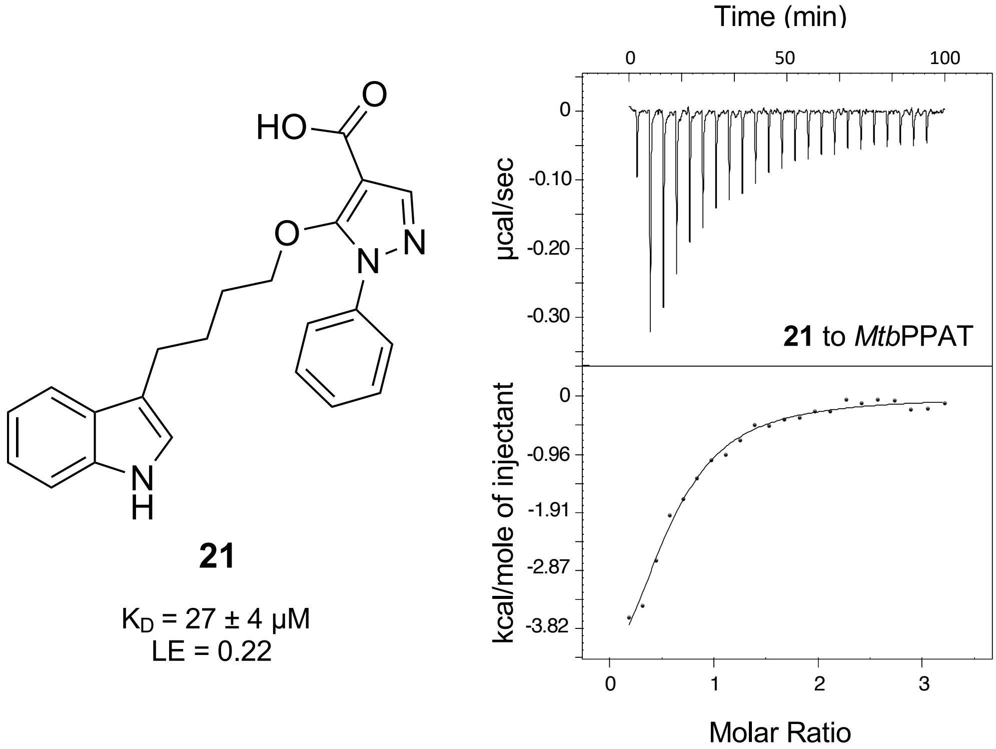

**21**

$K_D$ = 27 ± 4 µM
LE = 0.22

**Figure S11:** ITC analysis of *Mtb*PPAT binding to compound **21**. The upper panel shows the change in energy required to maintain a constant temperature during the titration of 100 µM *Mtb*PPAT (protomer concentration, equivalent to 17 µM hexamer) with compound **21** and the bottom panel shows the integrated heats of binding. Titrations were performed in the presence of 30 mM HEPES, pH 8.0, 20 mM NaCl, 0.5 mM TCEP and 5% (v/v) DMSO. To *Mtb*PPAT 22 × 2 µL compound **21** (1 mM) was injected. Data are fitted to a "one set of sites" binding model (solid line). A binding stoichiometry (*n*) of 0.5 (relative to the protomer) was determined. The data are representative of two independent experiments and the average $K_D$ determined ± range/2 is shown. The corresponding LE, in kcal.mol$^{-1}$.NHA$^{-1}$, is also shown.

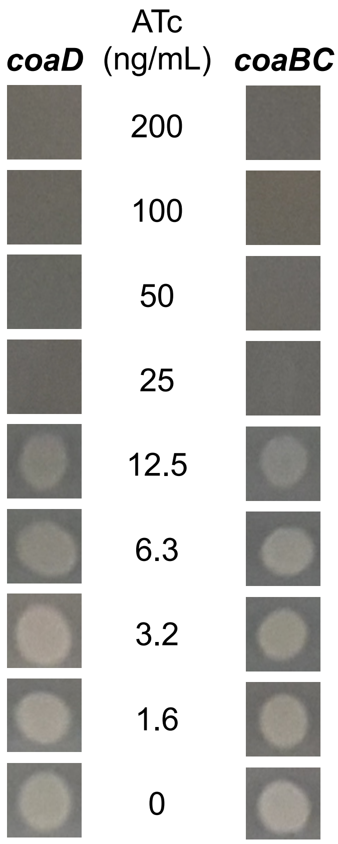

**Figure S12: ATc dose dependence of growth of *coaD* and *coaBC* CRISPRi *Mtb* mutants.** Strains were grown to early log phase before equivalent numbers of cells were spotted onto Middlebrook 7H10 agar containing kanamycin (25 µg/mL) and the indicated concentrations of ATc (ng/mL) and incubated at 37ºC for 7 days.

#### Supplementary Schemes

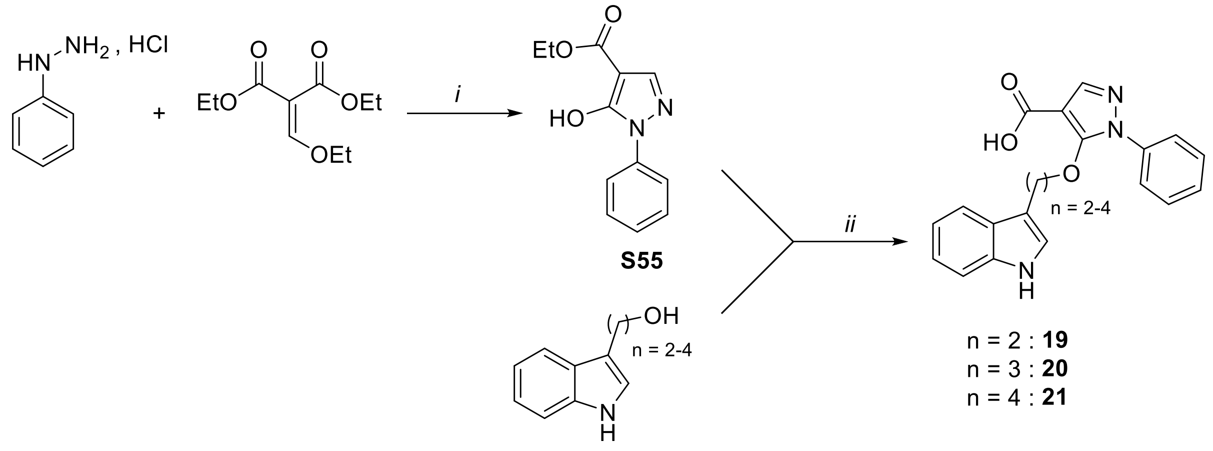

Reagents and conditions: (*i*) (a) $K_2CO_3$, $H_2O$, reflux, 2 h (b) 1 N HCl, 91%; *(ii)* (a) $PPh_3$, DEAD, THF, rt, 16 h (b) 1 N NaOH, EtOH, reflux, 2 h (c) 1 N HCl, 56-73% over 2 steps.

**Scheme S1:** Compounds **19** – **21** were synthesised *via* a Mitsunobu reaction between ethyl 5-hydroxy-1-phenyl-1*H*-pyrazole-4-carboxylate and the appropriate indole. This was followed by hydrolysis of the ethyl ester under alkaline conditions.

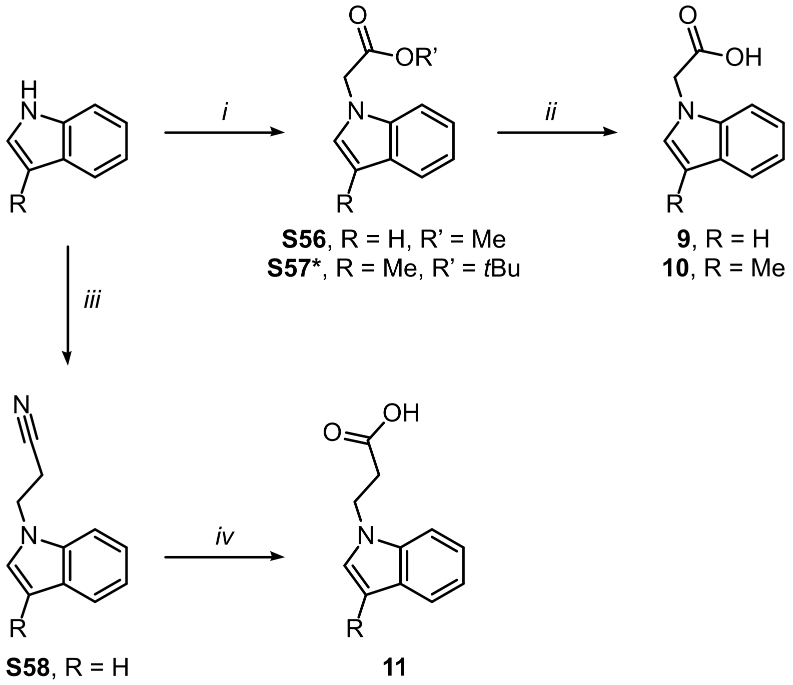

Reagents and conditions: (*i*) Methyl-2-bromoacetate, $K_2CO_3$, DMF, 16 h, rt, 20% (for **S56**), or *t*-butyl-2-bromoacetate, $K_2CO_3$, DMF, 24 h, rt (for **S57**, *not isolated prior to next step); (*ii*) THF, LiOH, MeOH, $H_2O$, 0˚C, 50 min, 98% (for **9**) or DCM, TFA, 0˚C then rt, 49% over 2 steps (for **10**); *(iii)* Benzyltrimethylammonium hydroxide, 1,4-dioxane, acrylonitrile, rt, 10 min, 68%; (*iv*) aq. KOH, reflux 3 h, 79%.

**Scheme S2:** Synthesis of analogues of fragment **2.** Compounds **9** and **10** were synthesised by alkylation of the relevant indole at the nitrogen using bromoacetic acid esters under basic conditions. Subsequent hydrolysis of the esters gave the desired fragments. Fragment **11** was synthesised via a propanenitrile intermediate; alkaline hydrolysis of the intermediate gave the desired fragment with a carboxylic acid moiety linked to the indole ring by two methylene units.

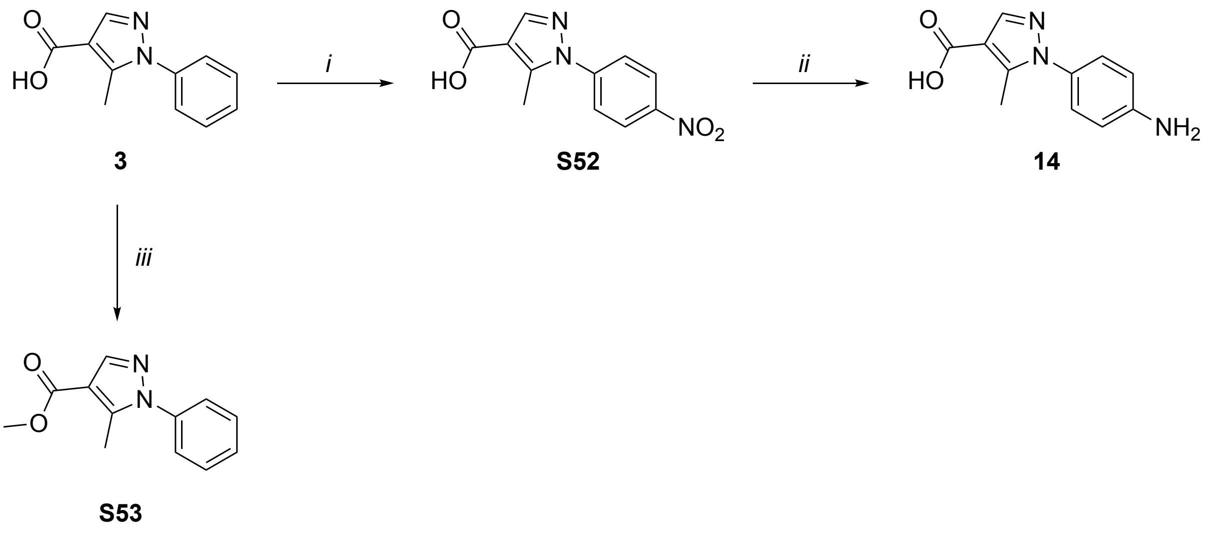

Reagents and conditions: (*i*) $H_2SO_4$, $HNO_3$, 0˚C, 10 min, 75%; (*ii*) $H_2$, Pd/C, MeOH, 12 h, 76%; (*iii*) (a) $PCl_5$, $Et_2O$, 0˚C, 24 h (b) MeOH, 86%.

**Scheme S3:** Synthesis of analogues of fragment **3.** Nitration of fragment **3** gave fragment **S52** and subsequent reduction of the nitro group of **S52** gave fragment **14**. Conversion of the carboxylic acid of fragment **3** to the methyl ester via an acid chloride intermediate gave fragment **S53**.

### Supplementary Table

**Table S1:** X-ray crystallography data collection and final refinement statistics.

| CMPD# | **3** | **6** | **11** | **12** | **20** |
|---|---|---|---|---|---|
| PDB ID | 6QMI | 6G6V | 6QMF | 6QMG | 6QMH |
| **Data collection*** | | | | | |
| X-ray source | SOLEIL PROXIMA -1 | DLS, I03 | DLS, I03 | DLS, I03 | DLS, I04-1 |
| Space group | *H*32 | *H*32 | *H*32 | *H*32 | *H*32 |
| Cell parameters: | | | | | |
| a = b [Å] | 97.99 | 97.75 | 97.75 | 98.09 | 98.96 |
| c [Å] | 113.96 | 114.71 | 112.73 | 113.09 | 115.51 |
| α=β=90°, γ=120° | | | | | |
| Resolution range [Å] | 34.03 – 1.78 (1.88 – 1.78) | 48.88 – 1.94 (2.05 – 1.94) | 67.78 – 1.77 (1.87 – 1.77) | 67.92 – 1.65 (1.74 – 1.65) | 49.48 – 1.84 (1.94 – 1.84) |
| No. of observations | | | | | |
| total | 168135 (24479) | 306649 (44359) | 398075 (58398) | 501122 (72856) | 256705 (33690) |
| unique | 20290 (2919) | 15776 (2262) | 20447 (2956) | 25454 (3690) | 19171 (2746) |
| $R_{merge}$ | 0.031 (0.734) | 0.075(1.487) | 0.059 (1.486) | 0.067 (1.589) | 0.099 (0.964) |
| I/σ(I) | 33.0 (2.8) | 24.7 (2.6) | 27.4 (2.7) | 26.4 (2.6) | 17.9 (2.5) |
| Completeness [%] | 99.9 (100.0) | 100.0 (100.0) | 100.0 (100.0) | 100.0 (100.0) | 100.0 (100.0) |
| Multiplicity | 8.3 (8.4) | 19.4 (19.6) | 19.5 (19.8) | 19.7 (19.7) | 13.4 (12.3) |
| **Refinement** | | | | | |
| Refinement program | PHENIX | PHENIX | PHENIX | PHENIX | PHENIX |
| Resolution [Å] | 34.03 – 1.78 | 48.87 – 1.94 | 67.78 – 1.77 | 67.92 – 1.65 | 49.48 – 1.84 |
| No. reflections | 20289 | 15776 | 20446 | 25454 | 19171 |
| $R_{work}/R_{free}$ [%] | 24.5/28.2 | 18.7/21.9 | 19.2/22.1 | 21.2/22.7 | 20.3/22.1 |
| RMS deviations | | | | | |
| Bonds [Å] | 0.008 | 0.007 | 0.007 | 0.007 | 0.008 |
| Angles [°] | 1.10 | 0.99 | 1.19 | 1.08 | 1.03 |
| Ramachandran | | | | | |
| Favoured [%] | 99.3 | 98.6 | 97.3 | 99.3 | 99.3 |
| Outliers [%] | 0 | 0 | 0 | 0 | 0 |

* Parameters shown in brackets are for the highest resolution shell

### Experimental Methods

##### *Expression and purification of* Mtb*PPAT*

The *MtbcoaD* gene was amplified by PCR from *Mtb* genomic DNA using a forward primer that incorporated a *Nde*I restriction site (5'-gccatatgacgggcgcggtatgcccaggg-3') and a reverse primer that incorporated a *Hind*III restriction site (5'-gcaagcttctaggtccgttcggtgttgagcctgtcccgc-3'). Following digestion with *Nde*I and *Hind*III, the PCR product was ligated to *Nde*I/*Hind*III-digested pET-28b. The resultant *MtbcoaD*-pET-28b plasmid encodes recombinant *Mtb*PPAT with a $His_6$-tag followed by a thrombin cleavage site (MGSSHHHHHHSSGLVPRGSH-) fused to the N-terminus of the protein. BL21(DE3) *E. coli* cells transformed with the *MtbcoaD*-pET-28b plasmid were grown at 37°C in Terrific Broth containing 50 µg/mL kanamycin, to an $OD_{600}$ of ~0.6. The temperature was then reduced to 20°C and isopropyl β-D-1-thiogalactopyranoside (IPTG) was added to a final concentration of 0.5 mM to induce protein expression. The expression was carried out for 20 h at 20°C and cells were harvested by centrifugation (10 min at 9000 × *g*). The cell pellets were resuspended in 30 mL of a lysis buffer consisting of 50 mM Tris-HCl, 500 mM NaCl, pH 7.8, with one dissolved EDTA-free protease inhibitor cocktail tablet (Roche) added, and were lysed at 4°C using an Avestin Emulsiflex High Pressure Homogeniser. Cell lysate was centrifuged at 35,000 × *g* for 30 min and the supernatant was loaded onto a Ni-NTA column (GE Healthcare) pre-equilibrated with the same lysis buffer. The column was washed sequentially using 100 mL of lysis buffer with 50 mM imidazole added, followed by 50 mL of lysis buffer with 100 mM imidazole added; the *Mtb*PPAT was subsequently eluted using a buffer consisting of 50 mM Tris-HCl, 500 mM NaCl, 1 M imidazole, pH 7.8. Elution and wash fractions were analysed by SDS-PAGE. Fractions containing the *Mtb*PPAT were pooled, concentrated using Vivaspin 20 centrifugal concentrators with a 10 kDa molecular weight cut-off (Vivaproducts) and injected onto a Superdex-200 HiLoad 26/60 gel filtration column (GE Healthcare) previously equilibrated in 30 mM HEPES, 200 mM NaCl, 5 mM $MgCl_2$, 0.5 mM TCEP, pH 8.0. Fractions were analysed by SDS-PAGE, and those containing the desired protein were pooled, divided into aliquots, flash frozen, and stored at −80 °C. Typically ~120 mg of *Mtb*PPAT was obtained from a 1 L culture.

##### *Fluorescence-based thermal shift experiments*

Fluorescence-based thermal shift experiments were performed in 96-well plates on a Roche LightCycler® 480. Each well contained 3 μM PPAT (protomer concentration, equivalent to 0.5 μM hexamer) in 50 mM HEPES, pH 7.2, containing 250 mM NaCl and 2.5 × SYPRO® Orange in a total volume of 100 μL per well. Fragments (which were dissolved to 100 mM in DMSO) were tested at a final concentration of 5 mM. All wells contained a final concentration of 5% (v/v) DMSO. Samples were heated from 37 to 85°C in the thermal cycler using a heating rate of 0.3°C/min. The thermal unfolding event was observed by excitation of the SYPRO® Orange dye at 490 nm; emission was detected at 530 nm. To generate melting curves, the relative fluorescence was plotted as a function of temperature. Melting temperatures ($T_m$ values, the maximum of the first derivative) were determined using the LightCycler® 480 software. Thermal shift ($\Delta T_m$) values were calculated by subtracting $T_m$ values measured for *Mtb*PPAT in the absence of fragment (but otherwise under identical conditions) from those measured in the presence of fragment.

##### *Ligand-based NMR binding experiments*

WaterLOGSY and STD experiments were performed on a Bruker Avance 700 Ultrashield spectrometer with Triple Resonance Inverse (TXI) cryoprobe (700 MHz) or a Bruker Avance 500 spectrometer with Triple Resonance Inverse (TCI) Automatic Tuning and Matching (ATM) cryoprobe (500 MHz). Samples (200 μL) were composed of 1 – 1.5 mM test compound with or without 20 μM *Mtb*PPAT (protomer concentration, equivalent to 3.3 μM hexamer) in 50 mM Tris-HCl, pH 7.8, 100 mM NaCl with 10% (v/v) $D_2O$ and 20 μM sodium 3-trimethylsilyltetradeuteriopropionate (TSP-$d_4$) as an internal standard. For competition experiments, ATP, CoA or the "reporter" fragment was included at a final concentration of 1 – 1.5 mM. The final concentration of DMSO-$d_6$ (the solvent in which fragments were solubilised) was ≤ 3% (v/v) and matched between samples in a given experiment. For NMR analysis, samples were transferred to 3-mm NMR capillary tubes (Hilgenberg GmbH) placed within 5-mm NMR tubes (Wilmad-LabGlass). WaterLOGSY and STD spectra were acquired at 278 K using an Autosampler and Icon-NMR Automation Suite, as described previously[1]. For WaterLOGSY experiments, 128 scans were acquired and for STD experiments, 256 scans were acquired. Data were processed using TopSpin 1.3 or 3.0 software.

##### *Isothermal titration calorimetry experiments*

ITC experiments were performed using a MicroCal iTC200 (Malvern) or Nano ITC (TA instruments). Titrations were performed at 25°C in 30 mM HEPES, pH 8.0 containing 200 mM NaCl, 5 mM $MgCl_2$, 0.5 mM TCEP, and 5% (v/v) DMSO or 30 mM HEPES, pH 8.0, 20 mM NaCl, 0.5 mM TCEP and 5% (v/v) DMSO, as indicated. Typically, 1 – 2 µL aliquots of the ligand solution were injected into 54 – 144 µM *Mtb*PPAT (protomer concentration, equivalent to 9 – 24 µM hexamer) with stirring at 1000 rpm (for titrations performed on the iTC200) or 300 rpm (for titrations performed on the Nano ITC). Data were fitted to a one set of sites independent binding model using Origin or NanoAnalyze software, following subtraction of the average heats measured in paired control titrations of ligand to buffer. As such small "heats of dilution" are difficult to measure absolutely, the value subtracted was optimised iteratively to minimize the $\chi^2$ calculated for the fit. As the direct titration of 144 µM *Mtb*PPAT (protomer concentration, equivalent to 24 µM hexamer) with fragments (at a concentration of 10 mM) produced low heats, fragment affinity was assessed indirectly by performing competition ITC experiments. In these experiments, 2 µL aliquots of CoA (600 µM) were injected into 54 µM *Mtb*PPAT (protomer concentration, equivalent to 9 µM hexamer) in the presence of 10 mM fragment. Fragment $K_D$ values were subsequently calculated from the apparent $K_D$ ($K_{D\ app}$) for coenzyme A (determined as described above) according to the method of Zhang and Zhang[2].

##### *X-ray crystallography*

Crystallisation of the *Mtb*PPAT apoprotein was performed at 293 K using the sitting-drop vapour diffusion method. Drops were composed of 2 µL of reservoir solution (15 – 30% 2-Methyl-2,4-pentanediol (MPD), 0.1 M Tris-HCl, pH 7.0 – 9.0, 10 – 30 mM $[Co(NH_3)_6]Cl_3$) and 2 – 3 µL of protein solution (20 mg/mL; stored in 25 mM Tris-HCl, pH 7.8, 150 mM NaCl). *Mtb*PPAT complexes were obtained by soaking compounds, dissolved in DMSO at concentrations of 10 – 100 mM, into crystals of the PPAT apoprotein in the previously set up drops (1:10 ratio, v/v) to reach a working concentration of 1 – 10 mM and left for 8 to 16 h. After this time crystals were fished out of the drop and flash frozen with liquid nitrogen using 50% MPD incorporated into the crystallisation mother liquor as a cryoprotectant.

X-ray datasets were collected on the in-house ICARUS diffractometer (X8 PROTEUM by Bruker AXS, Biochemistry Department, University of Cambridge) and at Diamond Light

Source synchrotron on beamlines I02, I03, I04, I04-1 and I24. *Mtb*PPAT crystals diffracted to between 1.5 and 2.5 Å resolution in H32 space group. The X-ray diffraction datasets were processed with autoPROC toolbox[3] using XDS[4] for indexing and SCALA/AIMLESS [5] for scaling. The *Mtb*PPAT structure was solved by molecular replacement using Phaser [6] with the 1TFU (PDB ID) structure as a search probe. Structure refinement was carried out by PHENIX program package[7] and Coot[8] was used for real-space refinement and further structure modifications. Small molecules (soaked compounds and natural ligands) were placed manually in Coot, guided by the observed Fo-Fc and 2Fo-Fc electron density and confirmed by omit maps after refinement was complete. Data collection and final refinement statistics are shown in **Table S1**.

**Mtb *strains and growth conditions***

*Mtb* H37RvMA[9] and its *coaD* and *coaBC* CRISPRi conditional knockdown derivatives (see below) were routinely grown in Difco Middlebrook 7H9 broth (BD) supplemented with Middlebrook albumin-dextrose-catalase (ADC) enrichment (BD), 0.2% (v/v) glycerol (Sigma-Aldrich) and 0.05% (v/v) Tween-80. Kanamycin was used at a final concentration of 25 µg/mL where required, and the anhydrotetracycline (ATc) inducer was used at concentrations up to 200 ng/mL in order to transcriptionally silence *coaD* and *coaBC* in the CRISPRi mutants.

***Construction of CRISPRi conditional knockdown mutants***

CRISPRi conditional knockdowns of *coaD* and *coaBC* were generated as previously described.[10] Briefly, a total of four and seven 20-24 bp sgRNA sequences with complementarity to the non-template target sequences of *coaD* and *coaBC*, respectively, were cloned into the CRISPRi plasmid (pLJR965) upstream of the dCas9 handle using golden gate cloning with *Bsm*BI. Following verification of the plasmids by Sanger sequencing, they were electroporated into *Mtb* H37RvMA[9] and the relative levels of ATc-induced growth repression by each sgRNA sequence was determined by spotting early log phase ($OD_{600}$ ~0.4) cultures on Difco Middlebrook 7H10 agar (BD) supplemented with Middlebrook oleic-albumin-dextrose-catalase (OADC) enrichment (BD), 0.5% (v/v) glycerol, kanamycin (25 µg/mL) and 2-fold serial dilutions of ATc ranging from 200 – 1.6 ng/mL. Since the 20 bp sgRNAs ATCGCGACTTCTTTGGCCAG and GAGGTGAACATGCGGGACCG produced the most stringent growth inhibition due to transcriptional silencing of *coaD* and *coaBC*, respectively (Fig. S12), these CRISPRi mutants were selected for use in all subsequent experiments.

***Effect of test compounds on* Mtb *growth***

The growth inhibitory effect of the test compounds was assessed using an Alamar Blue fluorescence-based assay, as previously described.[11] Briefly, 2-fold serial dilutions of test compounds in a 96-well microtiter plate were inoculated with *Mtb* at a cell density of $\sim 10^5$ CFU/mL. Plates were incubated at 37°C for 10 days, before 10 μL Alamar Blue solution was added and the plates were incubated for a further 24 h. Fluorescence (as an indication of growth) was measured using a SpectraMax i3x Multi-Mode Microplate Reader (Molecular Devices) in bottom-reading mode with excitation at 544 nm and emission at 590 nm.

***Synthetic Organic Chemistry***

**General Experimental Methods.** Solvents were distilled prior to use and dried by standard methods. Unless otherwise stated, $^1$H and $^{13}$C NMR spectra were obtained in $CD_2Cl_2$, $(CD_3)_2CO$, or $(CD_3)_2SO$ solutions using a Bruker 400 MHz AVANCE III HD Smart Probe, 400 MHz QNP cryoprobe, 500 MHz DCH cryoprobe spectrometer or Bruker 300 MHz AVANCE DRX spectrometer. Chemical shifts (δ) are given in ppm relative to the residual solvent peak, and the coupling constants (*J*) are reported in hertz (Hz).

Reactions were monitored by thin-layer chromatography (TLC) and liquid chromatography-mass spectrometry (LCMS) to determine consumption of starting materials. Flash column chromatography was performed using an Isolera Spektra One/Four purification system and the appropriately sized Biotage SNAP column containing KP-silica gel (50 μm). Solvents are reported as volume/volume eluent mixture where applicable.

LCMS was carried out using a Waters Acquity H-class Ultra Performance Liquid Chromatography (UPLC) system coupled to a Waters SQ Mass Spectrometer detector. Samples were separated using an Acquity UPLC HSS column operating at a flow rate of 0.8 mL/min. The eluent consisted of 0.1% formic acid in water (A) and acetonitrile (B); gradient, from 95% A to 5% A over a period of 4 or 7 min. Samples were detected at two wavelengths (254 and 280 nm) using a Waters Acquity TUV detector. All final compounds were of a purity greater than 95% by LCMS analysis unless otherwise stated. High resolution mass spectra (HRMS) were recorded using a Waters LCT Premier Time of Flight (TOF) mass spectrometer or a Micromass Quadrapole-Time of Flight (Q-TOF) spectrometer.

**Methyl 2-(1*H*-indol-1-yl)acetate (S56)**[12]

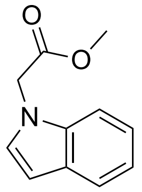

Potassium carbonate (3.53 g, 25.7 mmol) and methyl-2-bromoacetate (0.89 mL, 9.41 mmol) were added to a solution of indole (1.00 g, 8.55 mmol) in anhydrous DMF (32 mL). The reaction was stirred for 16 h at rt. Sodium bicarbonate (20 mL) was added and the product extracted in ethyl acetate (3 × 10 mL). The combined organic extracts were washed with water (3 × 50 mL) and brine (50 mL), dried ($Na_2SO_4$) and concentrated *in vacuo* to yield the ester **S56** (328 mg, 20%).

$^1$H NMR (500 MHz, $(CD_3)_2SO$): δ 7.54 (1H, d, *J* = 8.1), 7.36 (1H, d, *J* = 8.1), 7.32 (1H, d, *J* = 3.0), 7.11 (1H, t, *J* = 8.1), 7.02 (1H, t, *J* = 8.1), 6.45 (1H, d, *J* = 3.0), 5.12 (3H, s) and 4.92 (2H, s). $^{13}$C NMR (125 MHz, $(CD_3)_2SO$): δ 170.0, 136.8, 130.0, 128.5, 121.7, 120.8, 119.7, 110.2, 101.7, 52.5 and 47.3. HRMS (ESI$^+$): *m/z* calculated for $C_{11}H_{12}NO_2$ $[M + H]^+$ = 190.0863; found 190.0888.

**2-(1*H*-Indol-1-yl)acetic acid (9)**[12]

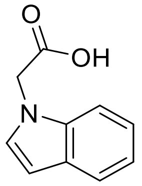

Methyl 2-(1*H*-indol-1-yl)acetate (0.468g, 2.48 mmol) was dissolved in THF (40 mL) and cooled to 0 °C. A solution of lithium hydroxide (0.416 g, 9.91 mmol) in methanol (10 mL) and water (10 mL) was added and the reaction was stirred at 0 °C for 50 min. Volatile solvents were removed *in vacuo* and the residue was acidified to pH 3 using aqueous HCl (1 M) and the product extracted in ethyl acetate (3 × 15 mL), dried ($Na_2SO_4$) and concentrated *in vacuo* to yield the carboxylic acid **9** (421 mg, 98%).

$^1$H NMR (500 MHz, $(CD_3)_2SO)$): δ 7.53 (1H, d, *J* = 8.2), 7.34 (1H, d, *J* = 8.2), 7.30 (1H, d, *J* = 3.0), 7.09 (1H, t, *J* = 8.2), 7.00 (1H, t, *J* = 8.2), 6.42 (1H, d, *J* = 3.0), and 4.92 (2H, s). $^{13}$C NMR (125 MHz, $(CD_3)_2SO)$): δ 170.9, 136.4, 129.8, 128.1, 121.1, 120.3, 119.1, 109.9, 100.7 and 47.7.

**2-(3-Methyl-1*H*-indol-1-yl)acetic acid (10)**

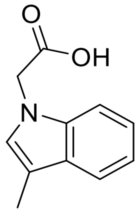

Potassium carbonate (3.51 g, 41.9 mmol) and *tert*-butyl-2-bromoacetate (1.3 mL, 9.5 mmol) were added to a solution of 3-methylindole (1.11 g, 8.47 mmol) in anhydrous DMF (30 mL). The reaction was stirred for 24 h at rt. Sodium bicarbonate (20 mL) was added and the product extracted in ethyl acetate (3 × 40 mL). The combined organic extracts were washed with water (3 × 100 mL) and brine (100 mL), dried ($Na_2SO_4$) and concentrated *in vacuo* to yield the intermediate *tert*-butyl ester (**S57**). The crude intermediate was dissolved in DCM (50 mL) and cooled to 0 °C, followed by dropwise addition of trifluoroacetic acid (1.3 mL, 17 mmol). The reaction was stirred at rt until completion, as determined by TLC and LCMS analysis, before the solvent was removed *in vacuo*. The residue was resuspended in an aqueous solution of sodium bicarbonate at pH 8 and washed with ethyl acetate (3 × 30 mL). The aqueous layer was then acidified to pH 3 using dilute aqueous HCl solution and the product extracted in ethyl acetate (3 × 60 mL), dried ($Na_2SO_4$) and concentrated *in vacuo* to give the carboxylic acid **10** (784 mg, 49% over two steps).

$^1$H NMR (400 MHz, $(CD_3)_2SO$): δ 12.95 (1H, s), 7.49 (1H, d, *J* = 7.8), 7.32 (1H, d, *J* = 7.8), 7.11 (1H, t, *J* = 7.8), 7.08 (1H, s), 7.02 (1H, t, *J* = 7.8), 4.93 (2H, s) and 2.25 (3H, s). $^{13}$C NMR (125 MHz, $(CD_3)_2SO$)): δ 170.8, 136.7, 128.4, 127.2, 121.2, 118.6, 118.5, 109.6, 109.3, 46.9 and 9.5. HRMS (ESI$^+$): *m/z* calculated for $C_{11}H_{12}NO_2$ $[M + H]^+$ = 190.0863; found 190.0909. Purity: 90%.

**3-(1*H*-Indol-1-yl)propanenitrile (S58)[13]**

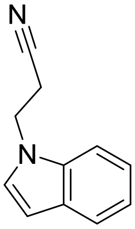

Benzyltrimethylammonium hydroxide (0.4 mL, 2.27 mmol) was dissolved in 1,4-dioxane (10 mL), followed by the addition of indole (1.00 g, 8.51 mmol) and acrylonitrile (1.0 mL, 15.3 mmol). The reaction was stirred at room temperature for 10 min, then neutralised with dilute acetic acid (1%, v/v, aqueous solution). The product was extracted in diethyl ether (3 × 20 mL), dried ($Na_2SO_4$) and concentrated *in vacuo*. Purification by flash column chromatography (gradient elution: 0-100% ethyl acetate in hexane) gave the propanenitrile (994 mg, 68%).

$^1$H NMR (400 MHz, $(CD_3)_2SO$): δ 7.60-7.54 (2H, m), 7.43 (1H, d, *J* = 3.7), 7.16 (1H, td, *J* = 7.3, 1.1), 7.05 (1H, td, *J* = 7.3, 1.1), 6.48 (1H, d, *J* = 3.7), 4.50 (2H, t, *J* = 6.5), 3.03 (2H, t, *J* = 6.5). $^{13}$C NMR (125 MHz, $(CD_3)_2SO$)): δ 135.6, 128.6, 128.3, 121.4, 120.6, 119.4, 119.0, 110.0, 101.4, 41.3 and 18.7. HRMS (ESI$^+$): *m/z* calculated for $C_{11}H_{11}N_2$ $[M + H]^+$ = 171.0917; found 171.0951.

**3-(1*H*-Indol-1-yl)propanoic acid (11)**

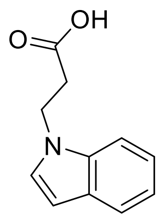

3-(1*H*-indol-1-yl)propanenitrile (0.450 g, 2.63 mmol) was dissolved in an aqueous solution of potassium hydroxide (10%, w/v, 10 mL) and heated to reflux for 3 h. The reaction mixture was cooled to rt and adjusted to pH 3 using acetic acid (10%, v/v). The product was extracted in ethyl acetate (3 × 20 mL), dried ($Na_2SO_4$) and concentrated *in vacuo* to give the carboxylic acid **11** (396 mg, 79%).

$^1$H NMR (400 MHz, $(CD_3)_2SO$): δ 12.36 (1H, s), 7.54 (1H, dd, J = 8.1, 1.1), 7.49 (1H, d, *J* = 8.1, 1.1), 7.35 (1H, d, *J* = 3.3), 7.13 (1H, td, *J* = 8.1, 1.1), 7.05 (1H, td, *J* = 7.3, 1.1), 6.48 (1H, d, *J* = 3.3), 4.40 (2H, t, *J* = 6.6) and 2.75 (2H, t, *J* = 6.6). $^{13}$C NMR (125 MHz, $(CD_3)_2SO$)): δ 172.6, 135.5, 128.7, 128.2, 121.1, 120.5, 119.1, 109.9, 100.9, 41.6 and 34.8. HRMS (ESI$^+$): *m/z* calculated for $C_{11}H_{12}NO_2$ $[M + H]^+$ = 190.0863; found 190.0890.

**5-Methyl-1-(4-nitrophenyl)-1*H*-pyrazole-4-carboxylic acid (S52)**

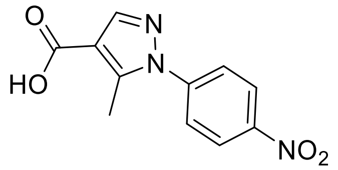

5-Methyl-1-phenyl-1*H*-pyrazole-4-carboxylic acid (51 mg, 0.250 mmol) was dissolved in concentrated sulphuric acid (0.3 mL). A mixture of concentrated nitric acid (70%, w/w, 0.1 mL) and concentrated sulphuric acid (0.3 mL) was added dropwise at 0 °C and the reaction mixture was stirred for 10 min. The solution was then poured onto a slurry of ice in water (5

g) and the resultant white precipitate collected by filtration, washed with ice cold water (2 × 5 mL) and dried *in vacuo* to give the nitrated product **S52** (46 mg, 75%).

$^{1}$H NMR (400 MHz, $(CD_3)_2SO$): δ 8.41 (2H, d, *J* = 9.6), 8.08 (1H, s), 7.90 (2H, d, *J* = 9.6) and 2.63 (3H, s). $^{13}$C NMR (125 MHz, $(CD_3)_2SO$)): δ 164.4, 146.6, 144.2, 143.6, 142.8, 125.9, 124.9, 114.2 and 11.9. HRMS (ESI$^{+}$): *m/z* calculated for $C_{11}H_{10}N_3O_4$ [M + H]$^{+}$ = 248.0671; found 248.0672. Purity: 90%.

**5-Methyl-1-(4-aminophenyl)-1*H*-pyrazole-4-carboxylic acid (14)**

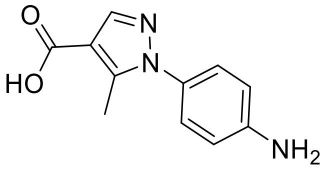

10% Palladium on carbon (5 mg) was added to a solution of 5-methyl-1-(4-nitrophenyl)-1*H*-pyrazole-4-carboxylic acid **S52** (20 mg, 0.081 mmol) in methanol (2.5 mL). The flask was sealed and flushed with hydrogen gas. The reaction was stirred under an atmosphere of hydrogen for 12 h, then filtered over Celite® and concentrated *in vacuo* to give aniline **14** (13 mg, 76%).

$^{1}$H NMR (400 MHz, $(CD_3)_2SO$)): δ 7.86 (1H, s), 7.13 (2H, d, *J* = 9.4), 6.70 (2H, d, *J* = 9.4) and 2.41 (3H, s). $^{13}$C NMR (125 MHz, $(CD_3)_2SO$)): δ 164.5, 143.8, 141.7, 140.1, 130.4, 127.3, 115.0, 113.0 and 12.3. HRMS (ESI$^{+}$): *m/z* calculated for $C_{11}H_{12}N_3O_2$ [M + H]$^{+}$ = 218.0930; found 218.0940.

**Methyl 5-methyl-1-phenyl-1*H*-pyrazole-4-carboxylate (S53)**

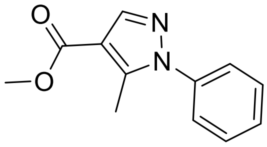

5-Methyl-1-phenyl-1*H*-pyrazole-4-carboxylic acid (50 mg, 0.250 mmol) was dissolved in diethyl ether (2.5 mL). Phosphorus pentachloride (200 mg, 0.375 mmol) was added at 0 °C and the reaction was stirred at room temperature for 24 h. The solvent was removed *in vacuo* and the residue resuspended in methanol (5 mL) at 0 °C. After stirring for 5 min, excess solvent was removed *in vacuo* to give the ester **S53** (46 mg, 86%).

$^{1}$H NMR (400 MHz, $(CD_3)_2SO$): δ 8.03 (1H, s), 7.60-7.50 (5H, m), 3.79 (3H, s), 2.52 (3H, s). $^{13}$C NMR (125 MHz, $(CD_3)_2SO$)): δ 163.4, 143.6, 141.3, 138.5, 129.4, 128.8, 125.4, 112.0,

51.2 and 11.7. HRMS (ESI$^+$): *m/z* calculated for $C_{12}H_{13}N_2O_2$ $[M + H]^+$ = 217.0972; found 217.0956. Purity: 92%.

**Ethyl 5-hydroxy-1-phenyl-1*H*-pyrazole-4-carboxylate (S55)**[14]

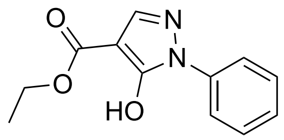

A solution of diethyl ethoxymethylenemalonate (2.38 g, 11.8 mmol), phenylhydrazine hydrochloride (1.71 g, 11.8 mmol) and $K_2CO_3$ (4.90 g, 35.4 mmol) in $H_2O$ was refluxed for 2 h. After cooling down to rt, the aqueous solution was washed with EtOAc and acidified to pH 2 (1 M HCl). The beige precipitate corresponding to ethyl 5-hydroxy-1-phenyl-1*H*-pyrazole-4-carboxylate was collected by filtration (2.50 g, 10.8 mmol, 91%).

$^1$H NMR (400 MHz, $(CD_3)_2CO$) δ 10.25 (s, 1H), 7.91 – 7.74 (m, 3H), 7.61 – 7.48 (m, 2H), 7.46 – 7.33 (m, 1H), 4.34 (q, *J* = 7.1 Hz, 2H), 1.35 (t, *J* = 7.1 Hz, 3H). LC-MS (ESI$^+$) *m/z* 233.2 $[M + H]^+$.

**5-(2-(1*H*-Indol-3-yl)ethoxy)-1-phenyl-1*H*-pyrazole-4-carboxylic acid (19)**

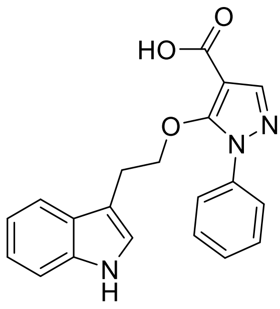

To a solution of tryptophol (0.139 g, 0.86 mmol), ethyl 5-hydroxy-1-phenyl-1*H*-pyrazole-4-carboxylate (0.200 g, 0.86 mmol) and $PPh_3$ (0.250 g, 0.95 mmol), in dry THF (7 mL), was added DEAD (0.150 mL, 0.95 mmol) at 0 °C, under nitrogen. The reaction mixture was stirred for 14 h at 20 °C and then concentrated *in vacuo*. The residue was purified by flash column chromatography (PE/EtOAc gradient 10−50%) to give pure ethyl 5-(2-(1*H*-indol-3-yl)ethoxy)-1-phenyl-1*H*-pyrazole-4-carboxylate intermediate as a pale yellow oil (267 mg, 0.71 mmol). $^1$H NMR (400 MHz, $CD_2Cl_2$) *δ* 8.12 (s, 1H), 7.92 (s, 1H), 7.63 – 7.56 (m, 2H), 7.54 (d, *J* = 7.9 Hz, 1H), 7.42 – 7.30 (m, 4H), 7.23 – 7.15 (m, 1H), 7.10 (ddd, *J* = 8.0, 7.1, 1.0 Hz, 1H), 6.98 (d, *J* = 2.4 Hz, 1H), 4.69 (t, *J* = 7.0 Hz, 2H), 4.32 (q, *J* = 7.1 Hz, 2H), 3.20 (td, *J* = 7.0, 0.8 Hz, 2H), 1.38 (t, *J* = 7.1 Hz, 3H).

To a solution of ethyl 5-(2-(1*H*-indol-3-yl)ethoxy)-1-phenyl-1*H*-pyrazole-4-carboxylate (0.112 g, 0.3 mmol) in EtOH (5 mL) was added 10% NaOH−$H_2O$ solution (5 mL). The reaction mixture was stirred for 14 h at rt. The reaction mixture was acidified with 1 M HCl solution, extracted with EtOAc, dried over $MgSO_4$, filtered, and concentrated *in vacuo*. The residue was then purified by flash column chromatography (DCM/MeOH gradient 1−10%) to give 5-(2-(1*H*-indol-3-yl)ethoxy)-1-phenyl-1*H*-pyrazole-4-carboxylic acid **20** as a beige solid (92 mg, 0.27 mmol, 74 % over 2 steps).

$^1$H NMR (400 MHz, $(CD_3)_2CO$) $\delta$ 10.01 (s, 1H), 7.93 (s, 1H), 7.63 (d, $J$ = 7.9 Hz, 2H), 7.55 (d, $J$ = 7.9 Hz, 1H), 7.45 – 7.31 (m, 4H), 7.15 – 7.06 (m, 2H), 7.05 – 6.95 (m, 1H), 4.78 (t, $J$ = 7.2 Hz, 2H), 3.21 (t, $J$ = 7.1 Hz, 2H). $^{13}$C NMR (125 MHz, $(CD_3)_2CO$) $\delta$ 163.8, 156.3, 143.1, 139.3, 130.1, 128.8, 128.6, 124.4, 124.3, 122.5, 119.9, 119.6, 112.6, 112.5, 111.5, 101.6, 77.1, 26.9. LC-MS (ESI$^+$) *m/z* 348.4 [M + H]$^+$. HRMS (ESI$^+$): *m/z* calculated for $C_{20}H_{17}N_3O_3Na$ [M + Na]$^+$ = 370.1162); found 370.1151.

**5-(3-(1*H*-indol-3-yl)propoxy)-1-phenyl-1*H*-pyrazole-4-carboxylic acid (20)**

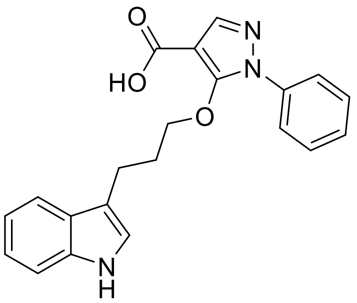

To a solution of 3-(3-hydroxypropyl)-1*H*-indole (0.150 g, 0.86 mmol), ethyl 5-hydroxy-1-phenyl-1*H*-pyrazole-4-carboxylate (0.200 g, 0.86 mmol) and $PPh_3$ (0.250 g, 0.95 mmol), in dry THF (7 mL), was added DEAD (0.150 mL, 0.95 mmol) at 0 °C, under nitrogen. The reaction mixture was stirred for 14 h at 20 °C and then concentrated *in vacuo*. The residue was purified by flash column chromatography (PE/EtOAc gradient 10−50%) to give ethyl 5-(3-(1*H*-indol-3-yl)propoxy)-1-phenyl-1*H*-pyrazole-4-carboxylate intermediate as a pale yellow oil (0.288 g, 0.74 mmol).

$^1$H NMR (400 MHz, $(CD_3)_2CO$) δ 9.89 (s, 1H), 7.98 (s, 1H), 7.83 – 7.73 (m, 2H), 7.52 (ddd, $J$ = 21.8, 10.7, 4.9 Hz, 3H), 7.45 – 7.36 (m, 2H), 7.16 – 7.09 (m, 1H), 7.08 – 7.01 (m, 1H), 6.98 (d, $J$ = 2.2 Hz, 1H), 4.54 (t, $J$ = 6.2 Hz, 2H), 4.30 (q, $J$ = 7.1 Hz, 2H), 2.81 (t, $J$ = 7.5 Hz, 2H), 2.17 – 2.07 (m, 2H), 1.33 (t, $J$ = 7.1 Hz, 3H).

To a solution of ethyl 5-(3-(1*H*-indol-3-yl)propoxy)-1-phenyl-1*H*-pyrazole-4-carboxylate (0.250 mg, 0.64 mmol) in EtOH (10 mL) was added 10% NaOH−$H_2O$ solution (10 mL). The reaction mixture was stirred for 14 h at rt. The reaction mixture was acidified with 1 M HCl solution, extracted with EtOAc, dried over $MgSO_4$, filtered, and concentrated *in vacuo*. The residue was then purified by flash column chromatography (DCM/MeOH gradient 1−10%) to give the desired compound as a beige solid (189 mg, 0.52 mmol, 70% over 2 steps).
$^1$H NMR (400 MHz, $(CD_3)_2CO$) δ 9.89 (s, 1H), 7.99 (s, 1H), 7.81 – 7.73 (m, 2H), 7.59 – 7.50 (m, 2H), 7.49 – 7.39 (m, 2H), 7.37 (dt, *J* = 8.2, 0.8 Hz, 1H), 7.10 (ddd, *J* = 8.2, 7.1, 1.1 Hz, 1H), 7.05 – 6.93 (m, 2H), 4.57 (t, *J* = 6.2 Hz, 2H), 2.79 (dd, *J* = 7.9, 7.2 Hz, 2H), 2.15 – 2.04 (m, 2H). $^{13}$C NMR (125 MHz, $(CD_3)_2CO$) δ 163.7, 156.4, 143.1, 139.4, 130.3, 128.9, 128.7, 124.7, 123.2, 123.1, 122.4, 119.7, 119.6, 115.4, 112.4, 101.6, 77.1, 31.5, 22.3. LC-MS (ESI$^+$) *m/z* 362.4 $(M + H)^+$. HRMS (ESI$^+$): *m/z* calculated $C_{21}H_{20}N_3O_3$ $[M + H]^+$ = 362.1499); found 362.1493.

**5-(3-(1*H*-indol-3-yl)propoxy)-1-phenyl-1*H*-pyrazole-4-carboxylic acid (21)**

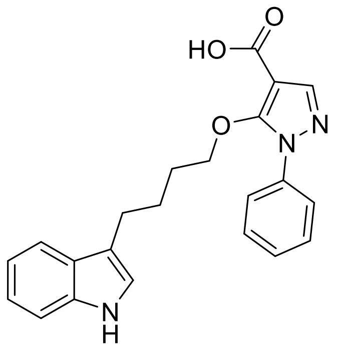

To a solution of 4-(1*H*-indol-3-yl)butan-1-ol (0.163 g, 0.86 mmol), ethyl 5-hydroxy-1-phenyl-1*H*-pyrazole-4-carboxylate (0.200 g, 0.86 mmol) and $PPh_3$ (0.250 g, 0.95 mmol), in dry THF (7 mL), was added DEAD (0.150 mL, 0.95 mmol) at 0 °C, under nitrogen. The reaction mixture was stirred for 14 h at 20 °C and then concentrated *in vacuo*. The residue was purified by flash column chromatography (PE/EtOAc gradient 10−50%) to give ethyl 5-(4-(1*H*-indol-3-yl)butoxy)-1-phenyl-1*H*-pyrazole-4-carboxylate intermediate as a pale yellow oil (0.254 g, 0.63 mmol). $^1$H NMR (300 MHz, $(CD_3)_2CO$) *δ* 9.92 (s, 1H), 7.89 (s, 1H), 7.71 – 7.67 (m, 2H), 7.52 (m, 3H), 7.45 – 7.36 (m, 2H), 7.11 – 6.97 (m, 3H), 4.49 (t, *J* = 6.0 Hz, 2H), 4.29 (q, *J* = 7.0 Hz, 2H), 2.74 (t, *J* = 7.5 Hz, 2H), 1.79 – 1.76 (m, 4H), 1.32 (t, *J* = 7.0 Hz, 3H).

To a solution of ethyl ethyl 5-(4-(1*H*-indol-3-yl)butoxy)-1-phenyl-1*H*-pyrazole-4-carboxylate (0.200 mg, 0.49 mmol) in EtOH (10 mL) was added 10% NaOH−$H_2O$ solution (10 mL). The reaction mixture was stirred for 14 h at rt. The reaction mixture was acidified with 1 M HCl solution, extracted with EtOAc, dried over $MgSO_4$, filtered, and concentrated *in vacuo*. The residue was then purified by flash column chromatography (DCM/MeOH gradient 1−10%) to give the desired compound as a beige solid (147 mg, 0.39 mmol, 56% over 2 steps).

$^1$H NMR (300 MHz, $(CD_3)_2CO$) δ 9.90 (s, 1H), 7.93 (s, 1H), 7.72 – 7.69 (m, 2H), 7.52 – 7.46 (m, 3H), 7.41 – 7.35 (m, 2H), 7.11 – 6.97 (m, 3H), 4.53 (t, *J* = 6.0 Hz, 2H), 2.73 (d, *J* = 7.5, 2H), 1.78 – 1.73 (m, 4H). $^{13}$C NMR (75 MHz, $(CD_3)_2CO$) δ 162.6, 155.1, 141.9, 138.0, 136.8, 128.9, 127.6, 127.5, 123.3, 122.8, 122.7, 121.1, 118.4, 118.3, 115.0, 111.2, 100.4, 76.0, 26.2, 24.4. LC-MS (ESI$^+$) *m/z* 375.4 $(M + H)^+$. HRMS (ESI$^+$): *m/z* calculated for $C_{22}H_{21}N_3O_3Na$ $[M + Na]^+$ = 398.1475); found 398.1480.

$^{1}H$ NMR Spectra (500 MHz, $(CD_3)_2SO$) of **9**

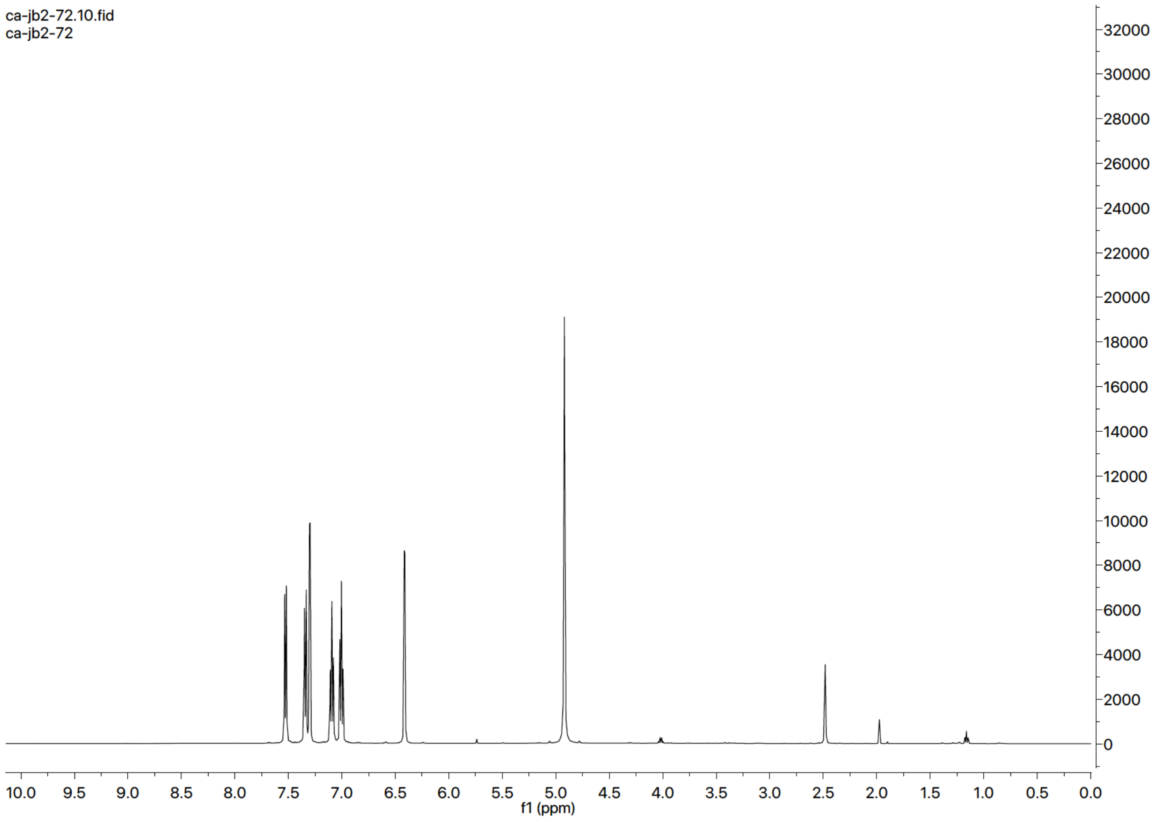

$^{13}C$ NMR Spectra (125 MHz, $(CD_3)_2SO$) of **9**

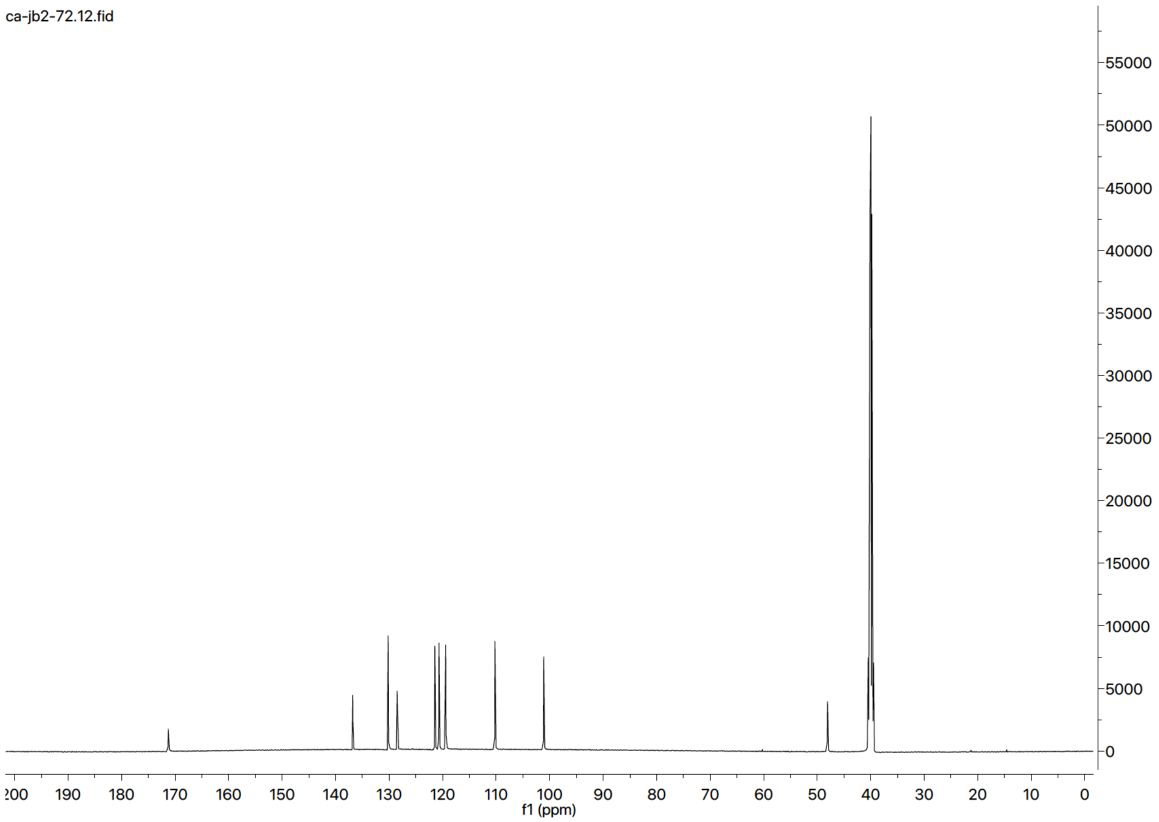

$^{1}H$ NMR Spectra (500 MHz, $(CD_3)_2SO$) of **10**

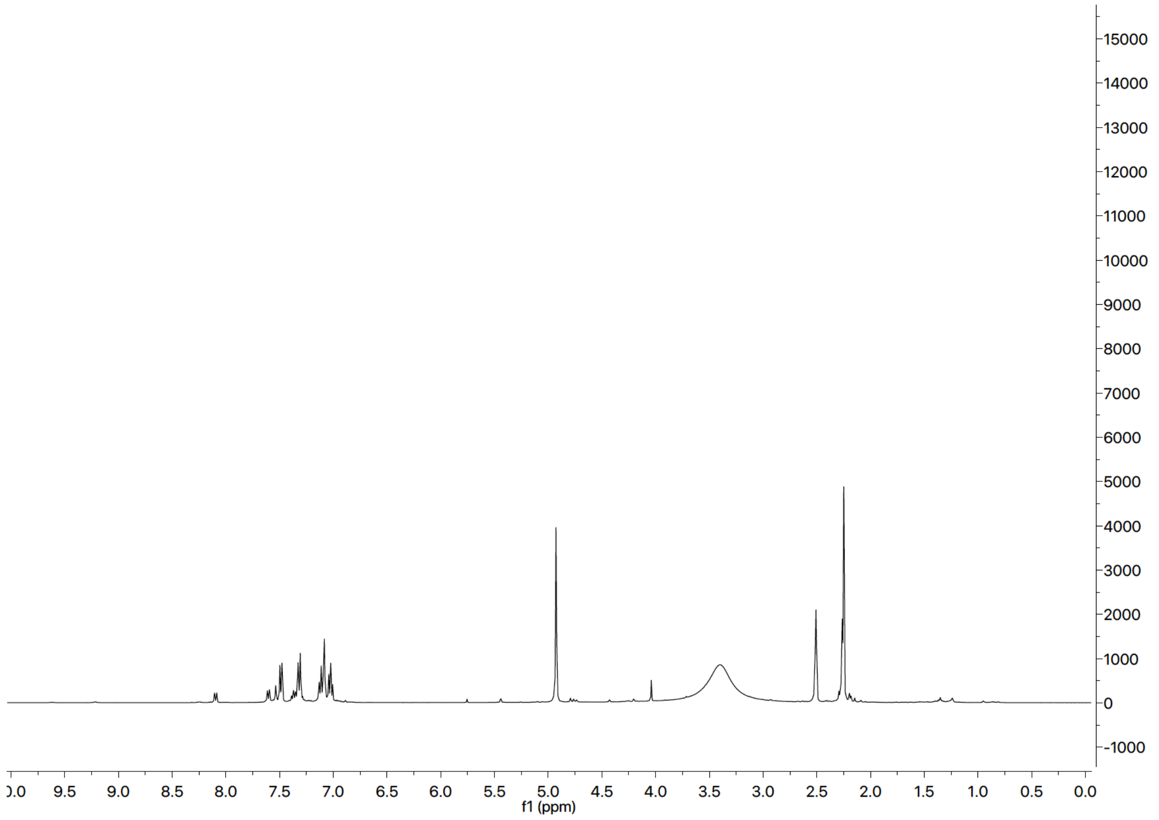

$^{13}C$ NMR Spectra (125 MHz, $(CD_3)_2SO$) of **10**

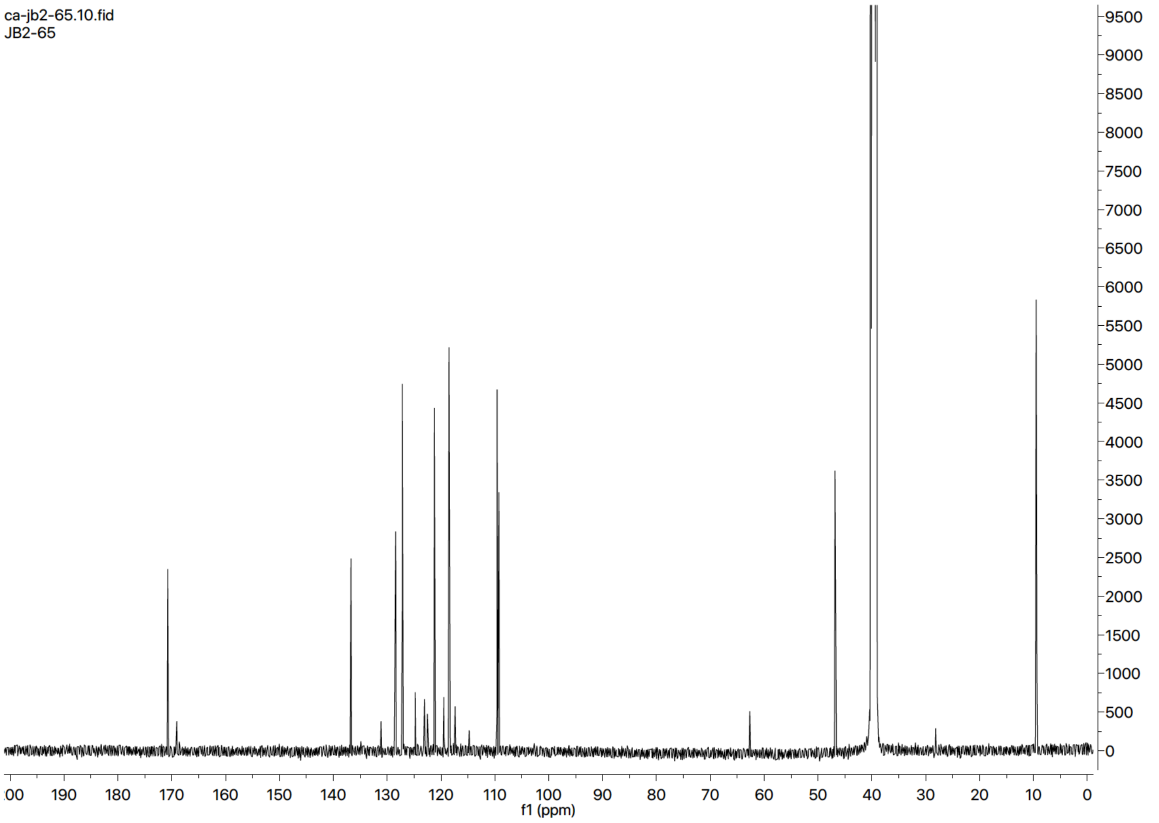

$^{1}H$ NMR Spectra (400 MHz, $(CD_3)_2SO$) of **S58**

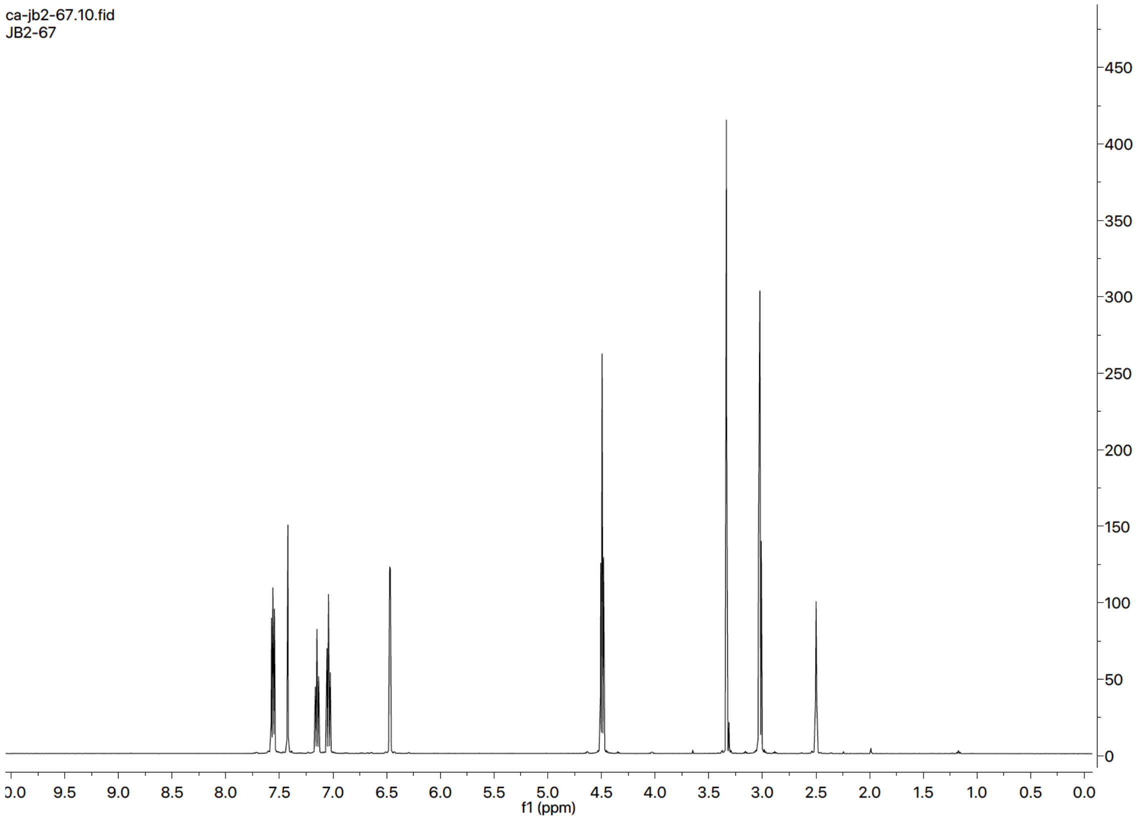

$^{13}C$ NMR Spectra (125 MHz, $(CD_3)_2SO$) of **S58**

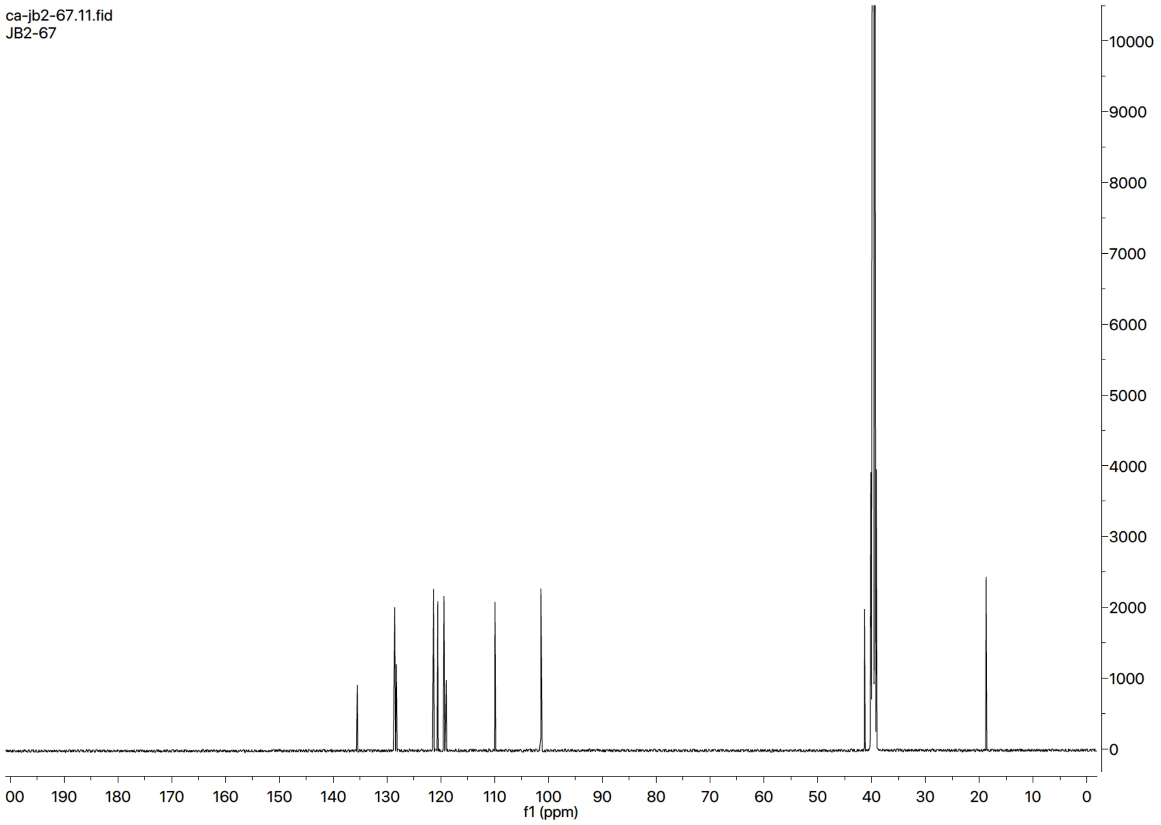

#### $^{1}$H NMR Spectra (400 MHz, $(CD_3)_2SO$) of **11**

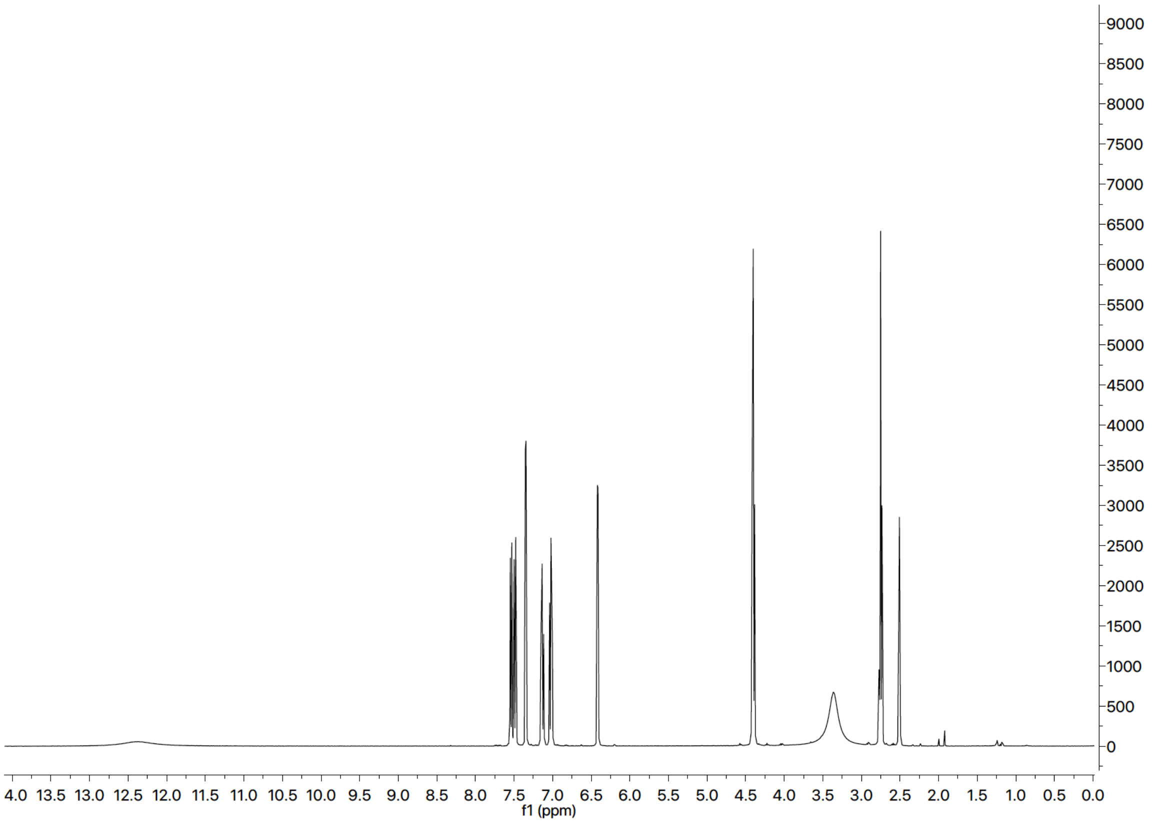

#### $^{13}$C NMR Spectra (125 MHz, $(CD_3)_2SO$) of **11**

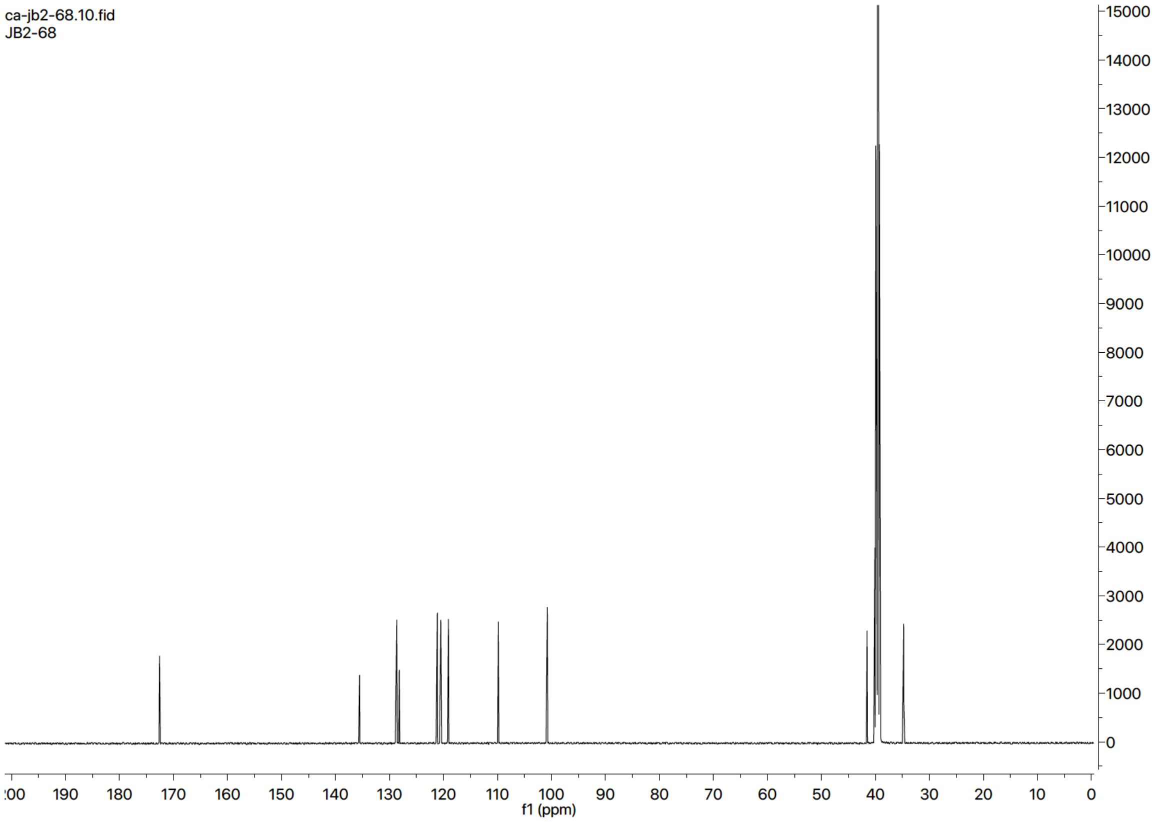

$^{1}H$ NMR Spectra (400 MHz, $(CD_3)_2SO$) of **S52**

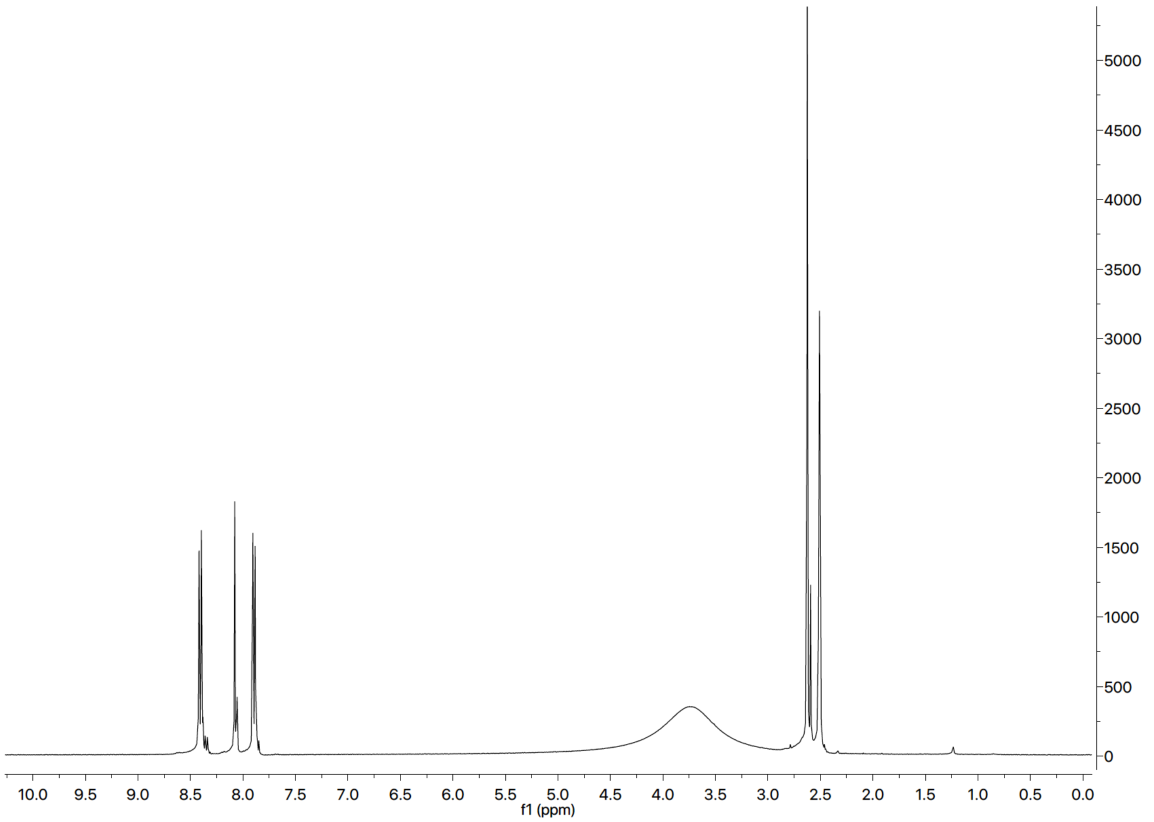

$^{13}C$ NMR Spectra (125 MHz, $(CD_3)_2SO$) of **S52**

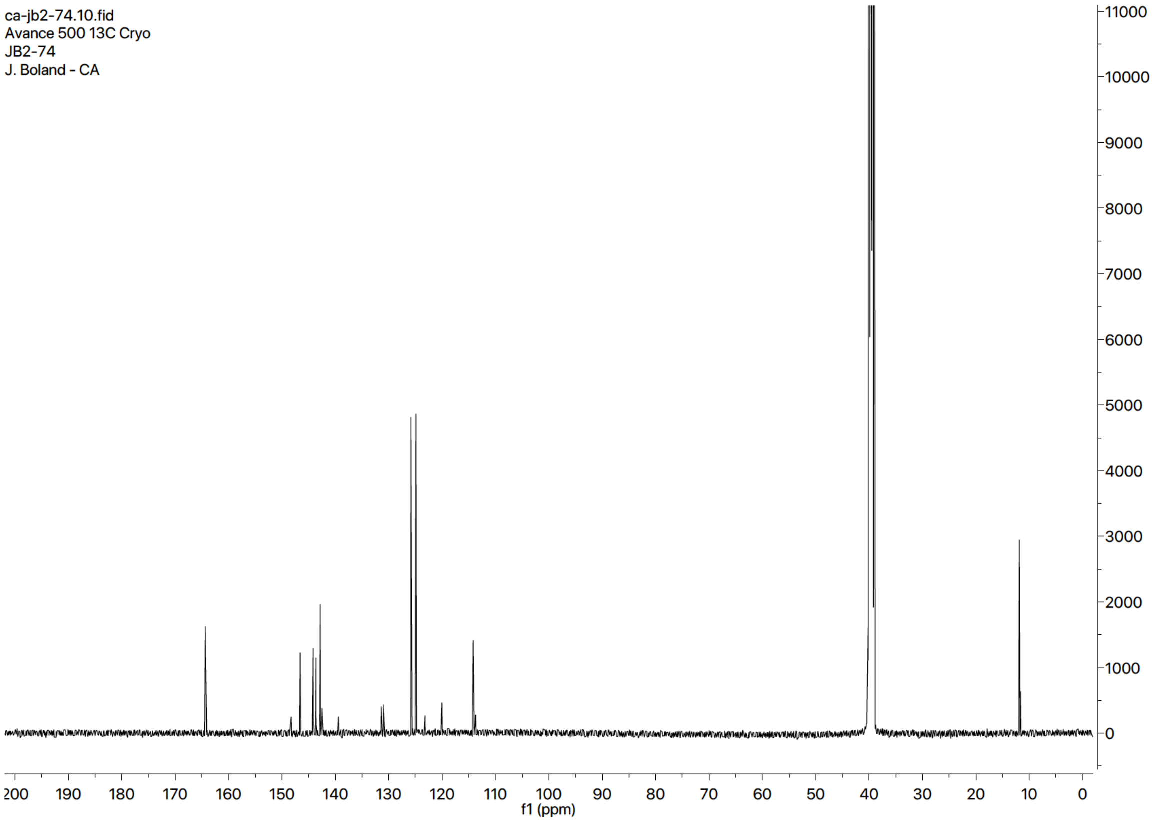

$^1H$ NMR Spectra (400 MHz, $(CD_3)_2SO$) of **14**

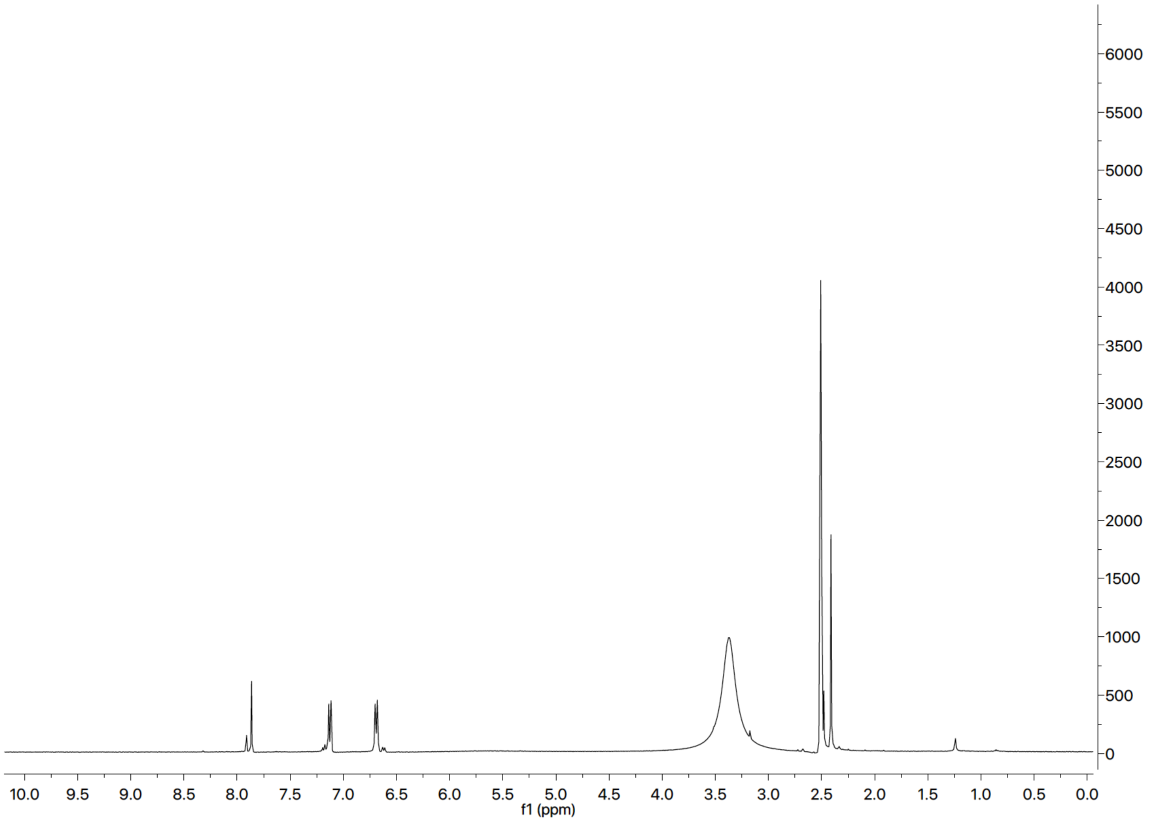

$^{13}C$ NMR Spectra (125 MHz, $(CD_3)_2SO$) of **14**

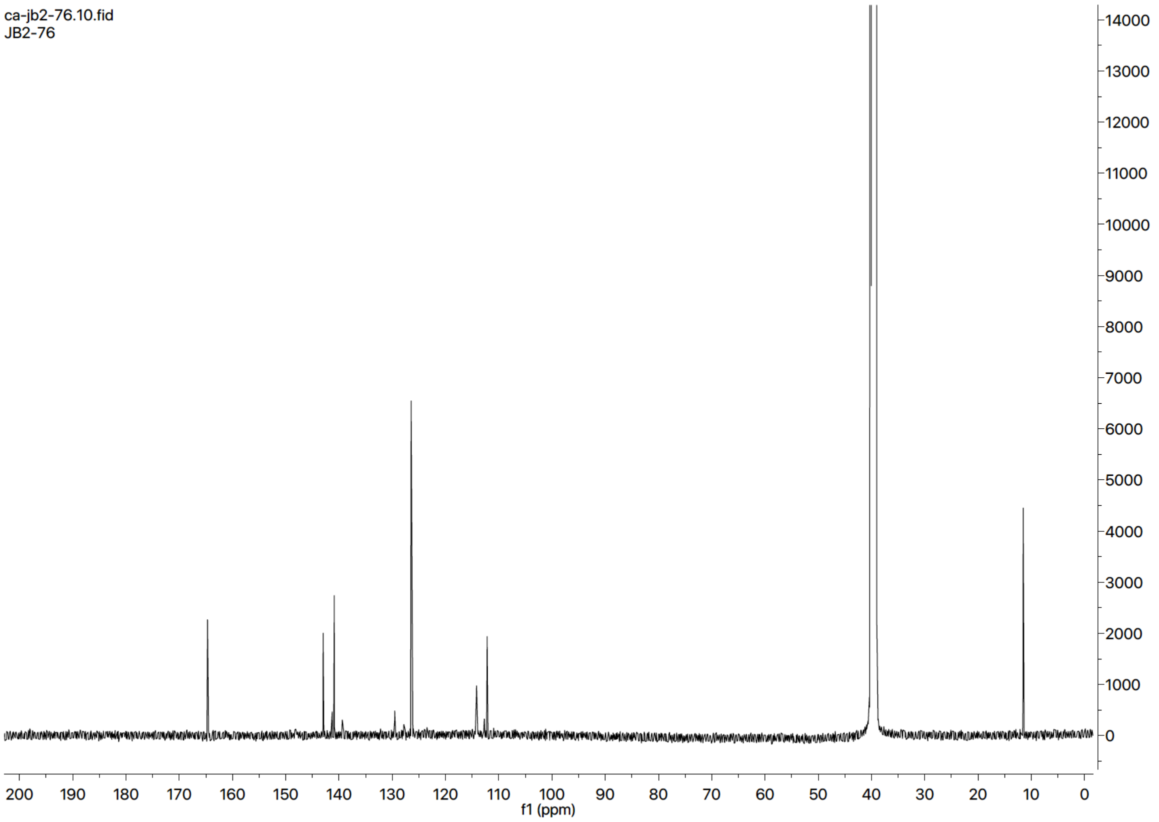

$^{1}H$ NMR Spectra (400 MHz, $(CD_3)_2SO$) of **S53**

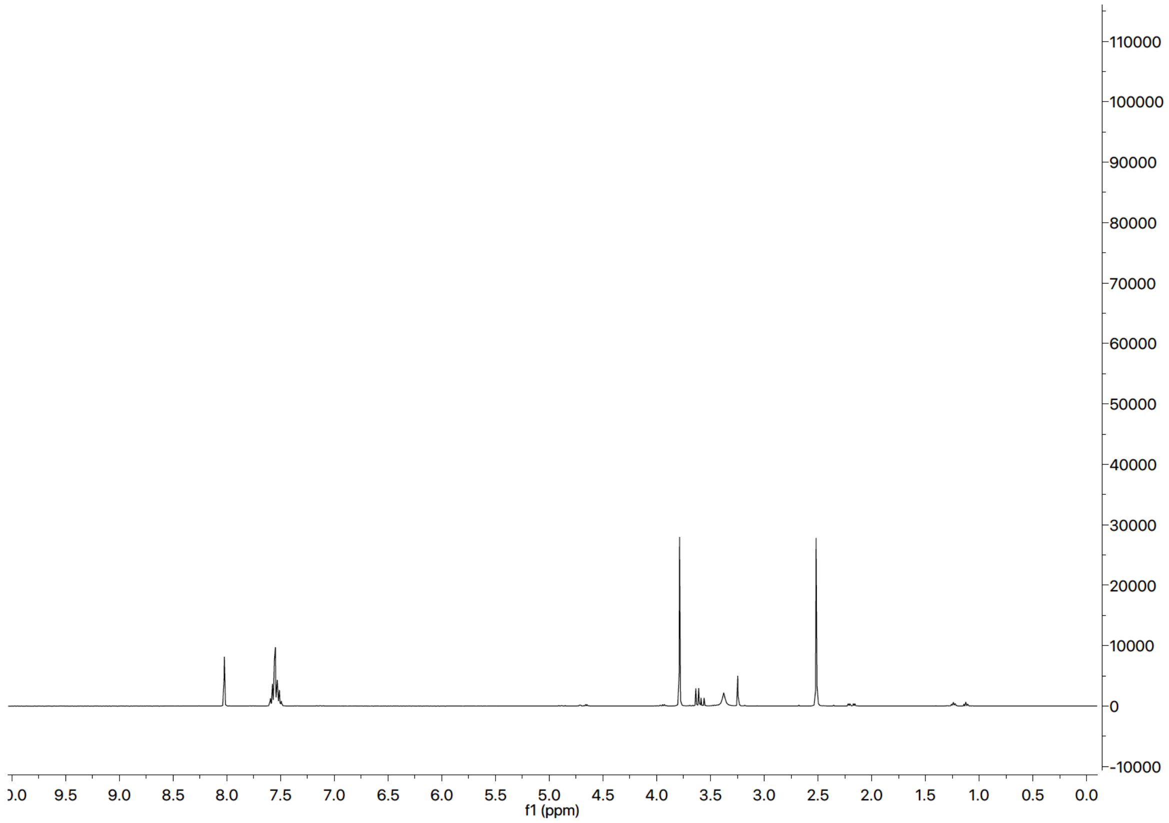

$^{13}C$ NMR Spectra (125 MHz, $(CD_3)_2SO$) of **S53**

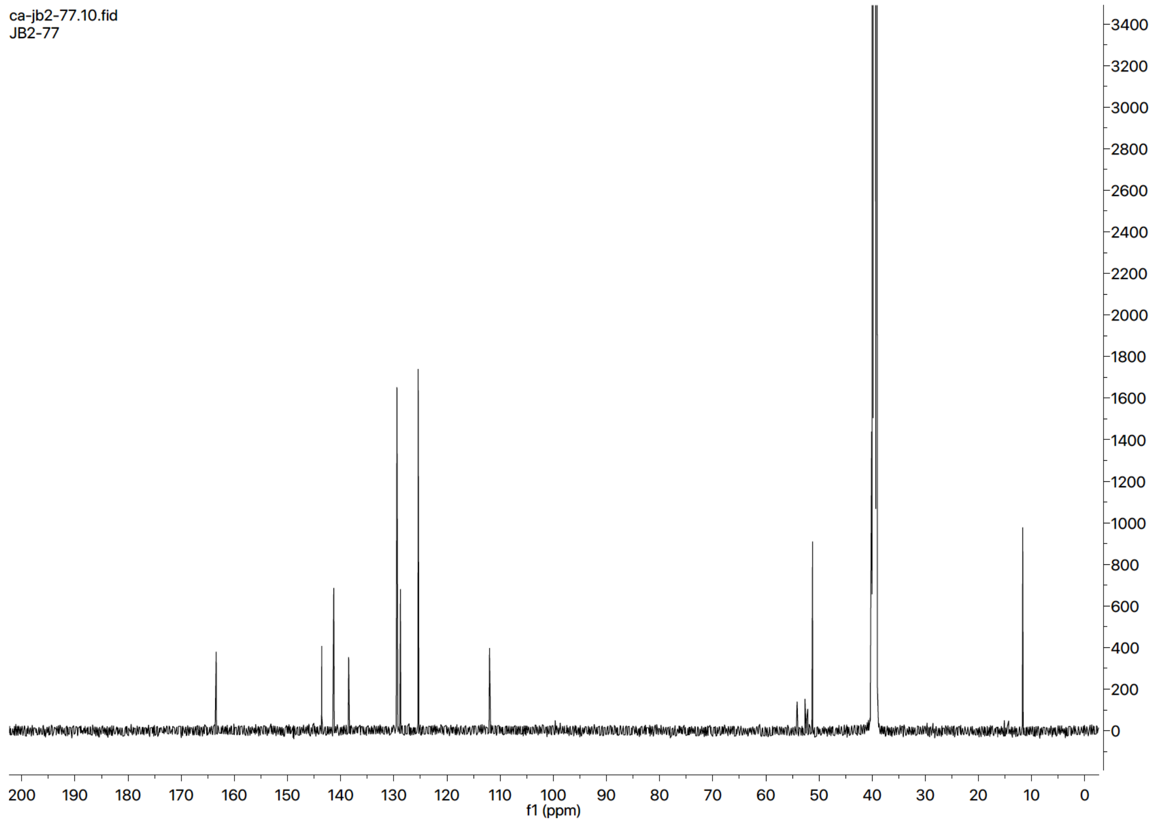

$^{1}H$ NMR Spectra (400 MHz, $(CD_3)_2CO$) of **19**

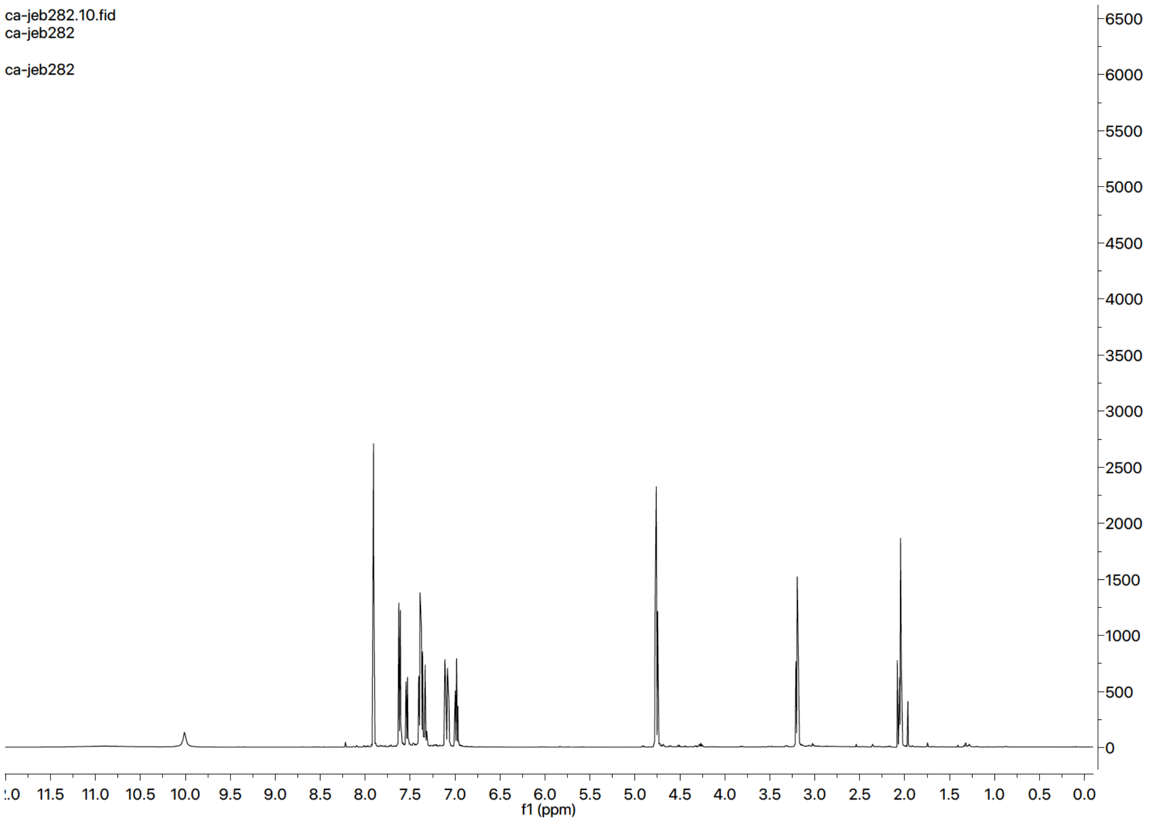

$^{13}C$ NMR Spectra (125 MHz, $(CD_3)_2CO$) of **19**

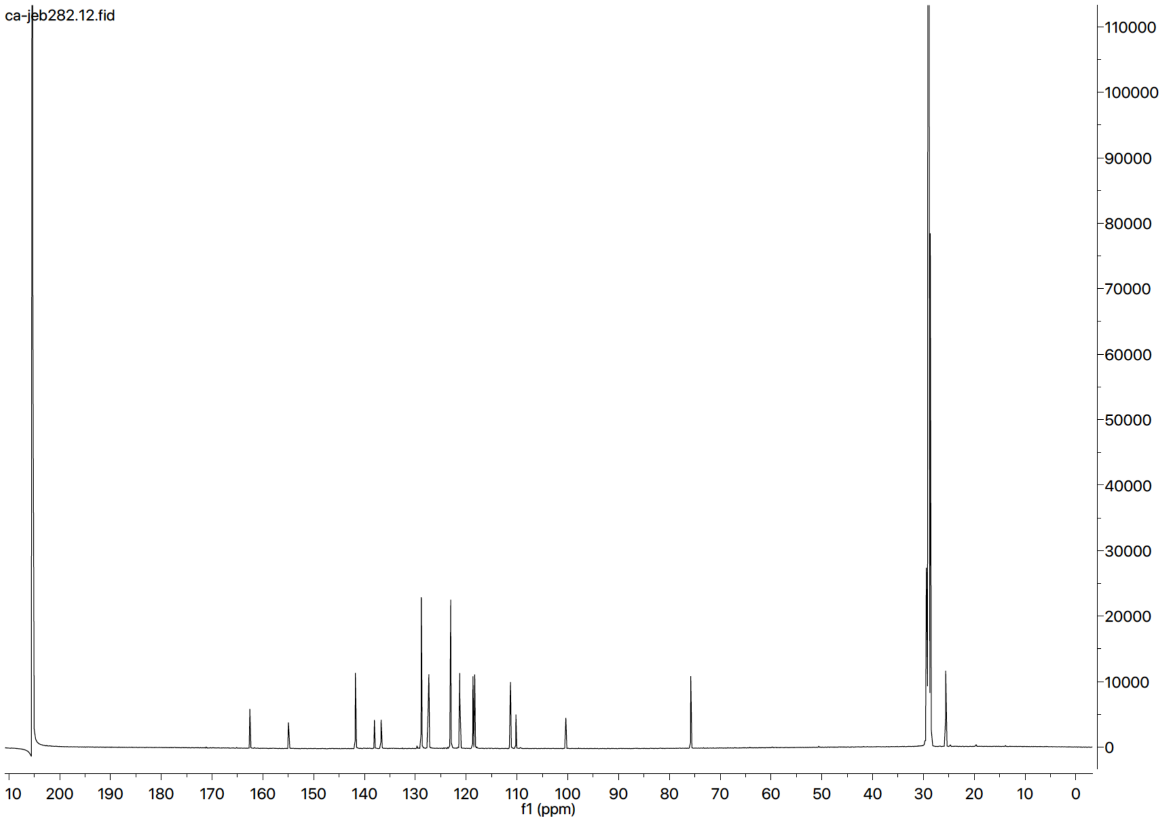

$^{1}$H NMR Spectra (400 MHz, $(CD_3)_2CO$) of **20**

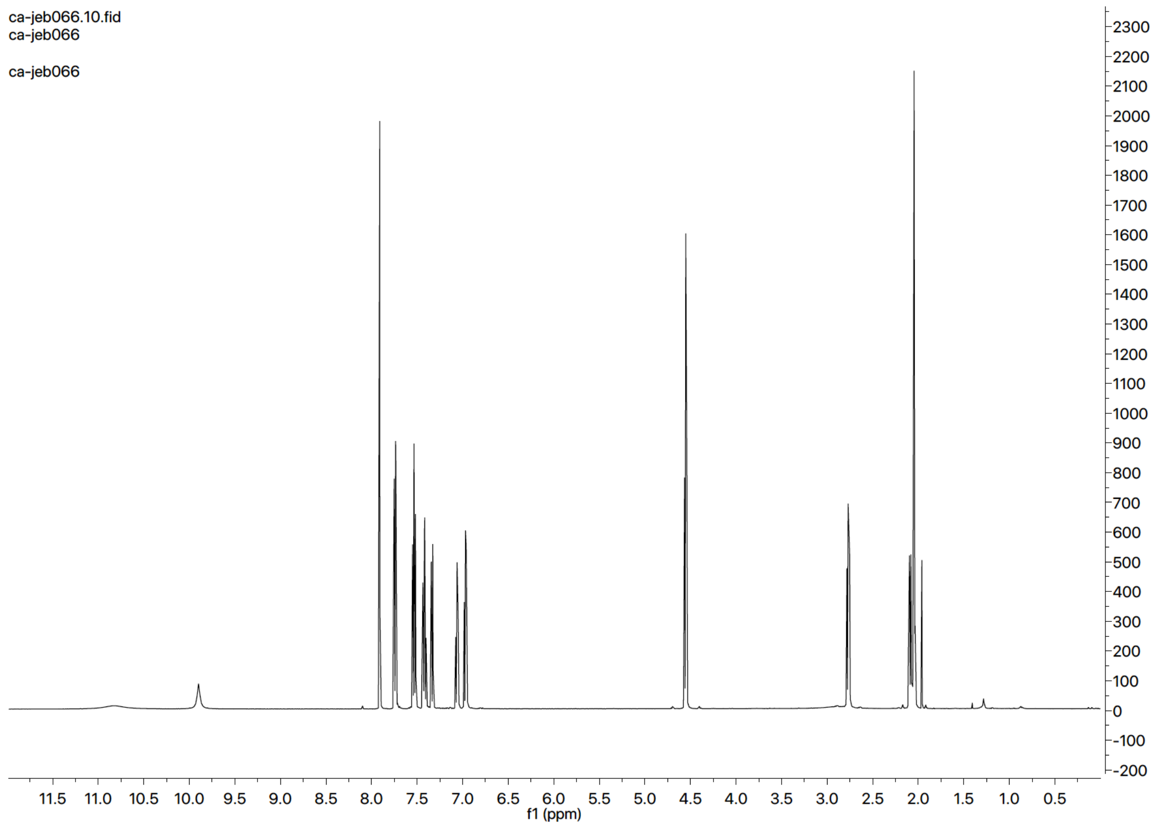

$^{13}$C NMR Spectra (125 MHz, $(CD_3)_2CO$) of **20**

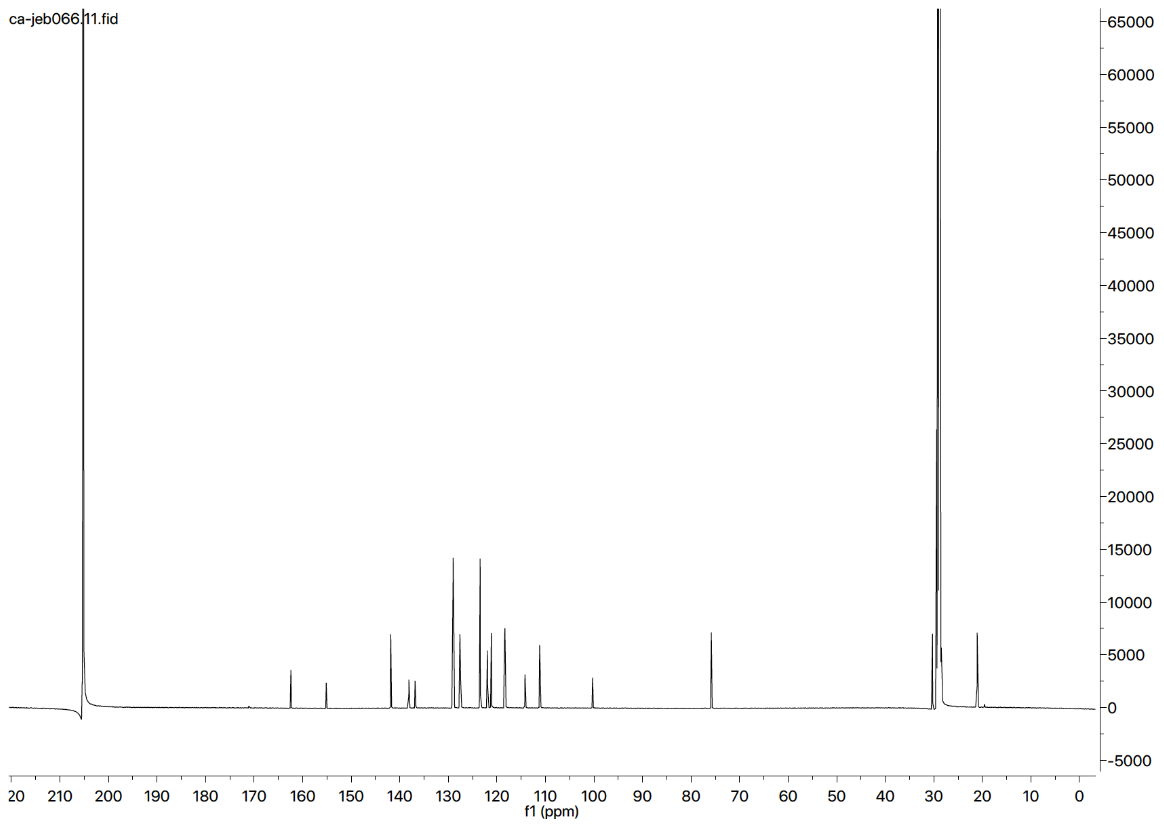

$^{1}H$ NMR Spectra (300 MHz, $(CD_3)_2CO$) of **21**

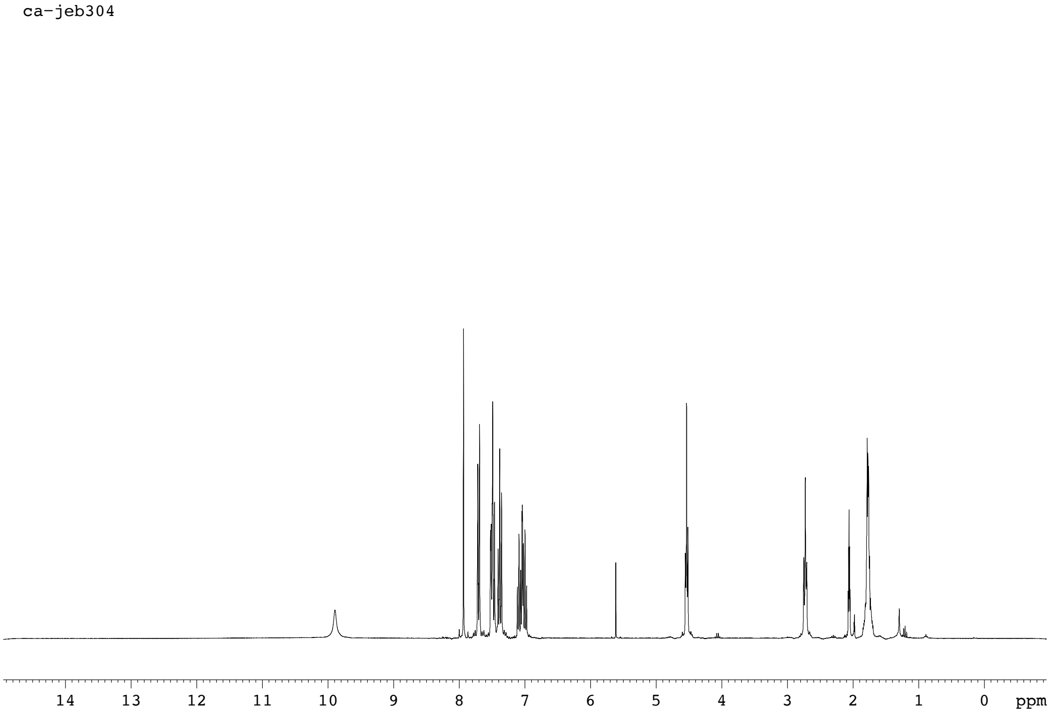

$^{13}C$ NMR Spectra (75 MHz, $(CD_3)_2CO$) of **21**

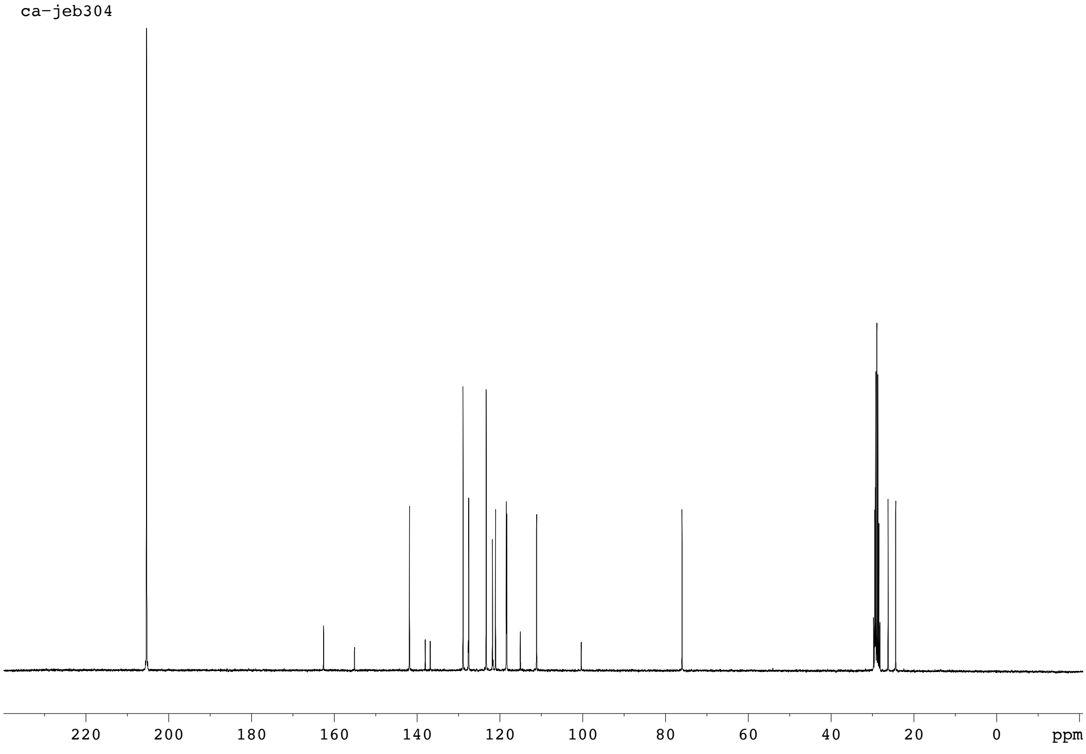
